# An automated, high throughput methodology optimized for quantitative cell-free mitochondrial and nuclear DNA isolation from plasma

**DOI:** 10.1101/2020.07.16.206987

**Authors:** Sarah A. Ware, Nikita Desai, Mabel Lopez, Daniel Leach, Yingze Zhang, Luca Giordano, S. Mehdi Nouraie, Martin Picard, Brett A. Kaufman

**Affiliations:** University of Pittsburgh School of Medicine, Department of Medicine, Division of Cardiology, Center for Metabolism and Mitochondrial Medicine, 200 Lothrop Street BST E1241, Pittsburgh, PA, 15261, USA; Optimize Laboratory Consultants, LLC, 104 Brunswick Place, Lansdale, PA,19446, USA; University of Pittsburgh School of Medicine, Department of Medicine, Division of Pulmonary, Allergy and Critical Care Medicine, 200 Lothrop Street, Pittsburgh, PA, 15261, USA; Columbia University Irving Medical Center, Departments of Psychiatry and Neurology, Division of Behavioral Medicine, 622 West 168^th^ Street, PH 1540 N, New York, NY, 10032, USA

**Keywords:** mitochondrial DNA (mtDNA), plasma, nucleic acid, DNA, liquid biopsy, cell-free, high throughput, liquid handling, magnetic particle processor, biomarker

## Abstract

Circulating, cell-free mitochondrial DNA (ccf-mtDNA) and nuclear DNA (ccf-nDNA) are under investigation as biomarkers for various diseases. Optimal ccf-mtDNA isolation parameters, like those outlined for ccf-nDNA, have not been established. Here, we optimized a protocol for both ccf-mtDNA and ccf-nDNA recovery using a magnetic bead-based isolation process on an automated 96-well platform. Using the optimized protocol, our data show 6-fold improved yields of ccf-mtDNA when compared to the starting protocol. Digestion conditions, liquid handling characteristics, and magnetic particle processor programming all contributed to increased recovery and improved reproducibility. To our knowledge, this is the first high-throughput approach optimized for mtDNA and nDNA recovery and serves as an important starting point for clinical studies.

**Graphical Abstract:** 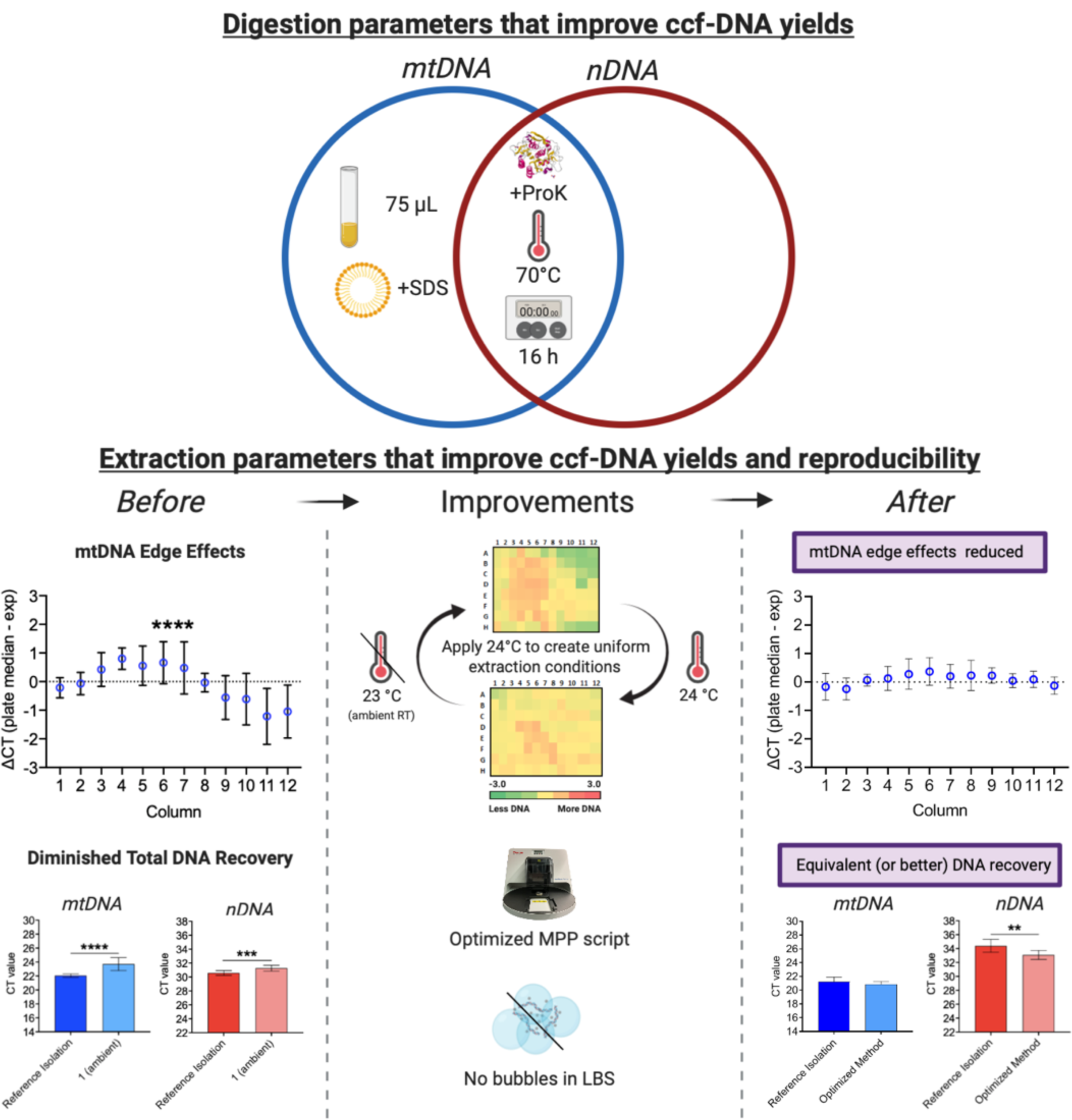

## Introduction

The presence of mitochondrial DNA (mtDNA) in plasma and serum was first documented more than 20 years ago (1). Nearly 10 years later, an association between elevated levels of circulating, cell-free mtDNA (ccf-mtDNA) and higher rates of mortality in trauma patients was established (2–6). Now, at an accelerating rate, levels of ccf-mtDNA are being examined in a variety of human disease states including HIV infections (7), major depressive disorder (8), psychological distress (9), type 2 diabetes (10), and myocardial ischemia-reperfusion injury (11, 12). Before we can understand the prevalence of elevated ccf-mtDNA levels in disease, a robust method to isolate ccf-DNA from large sample sizes is needed.

Surprisingly, while the discovery of ccf-mtDNA is relatively recent, the presence of nuclear DNA (nDNA) in the circulation has been known for over 70 years (13). In fact, numerous associative studies have since identified differences among the characteristics of ccf-nDNA (more generally termed ccf-DNA), between healthy individuals and diseased patients (14–20). Lately, ccf-DNA has been evaluated for its utility as a reliable diagnostic and prognostic biomarker. Such investigations have demonstrated an association between increased levels of ccf-DNA, worsened disease states, and poor clinical outcomes (8, 21, 22) for various conditions, including cancers (14, 23–25), chronic obstructive pulmonary disease exacerbations (21), and acute myocardial infarction (26). However, there has been a lack of consensus between protocols for ccf-DNA analysis, including blood draws, sample processing, and isolated ccf-DNA storage. This preparation inconsistency prevents the use of ccf-DNA analysis in routine clinical practice. To address this issue, a literature review identified preanalytical parameters that affected ccf-DNA concentration and fragmentation (27), including collection tube selection, processing time, and storage conditions.

Increasingly, studies seek to determine whether ccf-mtDNA or ccf-nDNA differentially associate with pathology. However, all prior procedural studies were not designed to track and optimize recovery of mtDNA and nDNA separately. mtDNA is a small circular genome up to 4 orders of magnitude smaller than linear nDNA fragments, and both genomes exhibit substantial differences in their native nucleoprotein complexes that could affect their isolation (28–30). In this study, we sought to identify an isolation method that was amenable to high-throughput platforms without sacrificing maximal, quantitative recovery of both ccf-mtDNA and ccf-nDNA. Various technologies have been used to isolate ccf-DNA, including column- and <u>m</u>agnetic <u>b</u>ead (MB)-based DNA affinity approaches, as well as alcohol-based precipitation methods. For large sample numbers, the column-based method has been the most commonly used, as precipitation methods that involve multiple centrifugation steps create bottlenecks in the workflow. Several reports have established that MB-based isolation methods yield high-quality DNA in similar quantities to column-based methods (25, 31–34). It has also been reported that MB-based extractions were the only method able to isolate key sequences when compared to other precipitation- and column-based methods (31). Based on this characteristic, we selected a commercial MB protocol for ccf-mtDNA isolation.

In this study, we focused on the maximization of yield and minimization of variability during ccf-DNA isolation from human plasma. We optimized the digestion and extraction steps by measuring both ccf-mtDNA and ccf-nDNA (total ccf-DNA) recovery, while focusing on yield, throughput, and reproducibility. For wide-scale screening of patient samples, any DNA isolation method should be adapted to high throughput platforms to reduce processing time without introducing variability. As such, we report crucial parameters that influence quantitative recovery of ccf-mtDNA and ccf-nDNA using a high throughput, magnetic particle separator-based workflow.

## Results

There are two aims of this study. The first aim is to identify chemical and laboratory parameters of plasma digestion and ccf-DNA extraction component steps that influence yields and provide evidence-based recommendations for such parameters that result in maximal ccf-DNA recovery. The first section, titled “*Summary of ccf-DNA isolation and quantification”*, introduces the basic DNA isolation process. “*Optimization of digestion parameters*” addresses optimal digestion conditions while “*Optimization of extraction parameters”* addresses conditions for ccf-DNA extraction. The second aim is to identify technical aspects of the ccf-DNA isolation process that limit its reproducibility for high throughput experiments. To that end, “*Identifying the source of variation in 96-well plate experiments”* identifies variability in 96 technical replicates across a 96-well plate while the remaining two sections address modification of magnetic particle processor (MPP) programming and liquid handling characteristics that improve both reproducibility and ccf-DNA yields.

### Summary of ccf-DNA isolation and quantification

The process of isolating and quantifying ccf-DNA from plasma required plasma digestion, ccf-DNA extraction, and qPCR (Fig. 1). As a brief overview, plasma was first incubated with Proteinase K (ProK) and sodium dodecyl sulfate (SDS) at 70 °C for 16 h to digest and disrupt protein and lipid structures, such as extracellular vesicles that may compartmentalize DNA and interfere with the interaction with magnetic beads (MB). Next, MB and Lysis/Binding Solution (LBS) were added to the digested plasma. The DNA-bound MB were transferred by the MPP into each subsequent solution. These steps removed protein contaminants (Wash 1), condensed and washed the DNA (Washes 2 and 3), then eluted DNA in Elution Solution (ES). DNA was stored at -20 °C before ccf-mtDNA and ccf-nDNA abundance were simultaneously measured by duplex qPCR.

**Figure 1.**
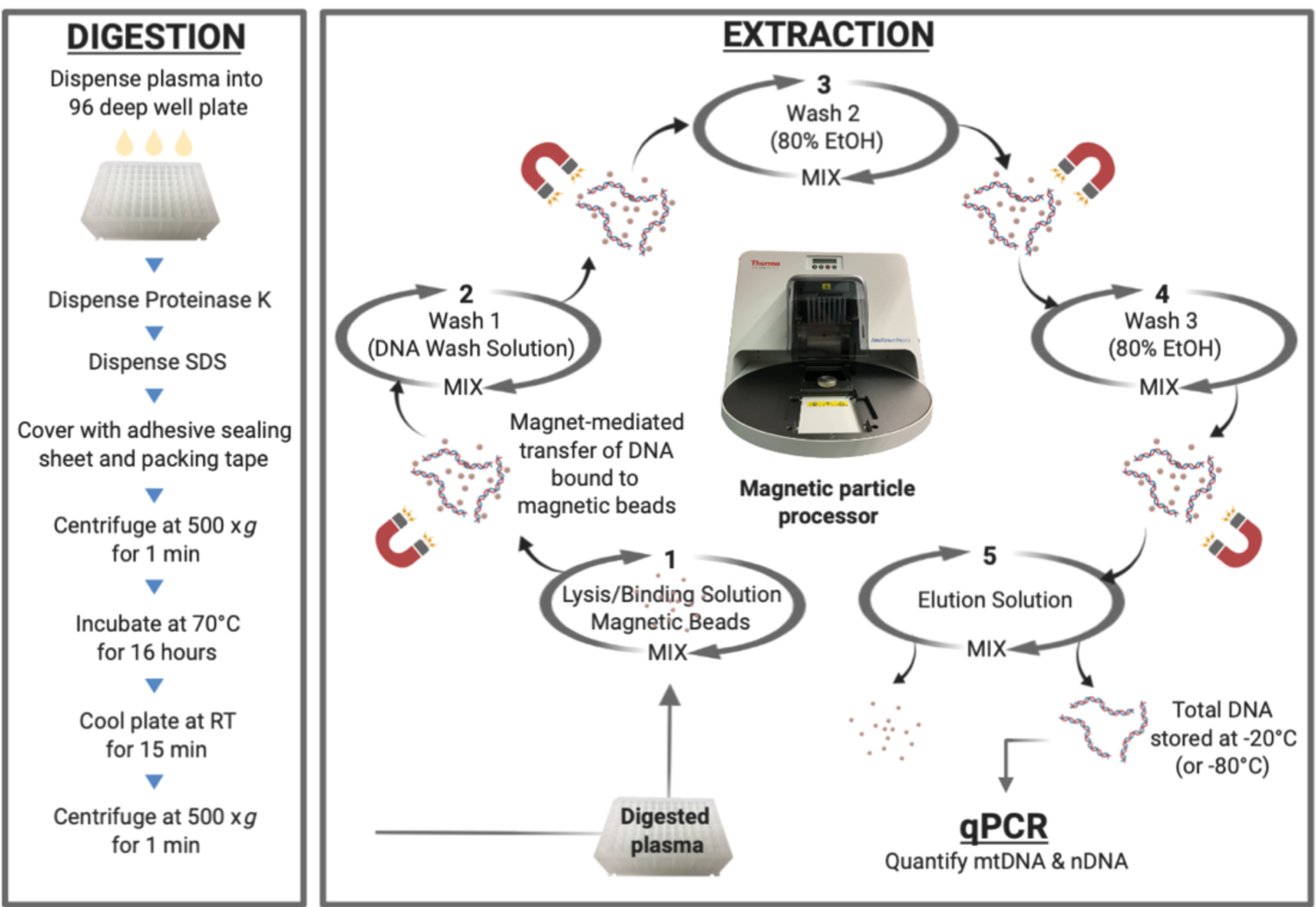
Workflow of automated isolation of ccf-DNA from plasma. Plasma, Proteinase K, and SDS were dispensed into a deep well plate and incubated at 70 °C for 16 hours (h) to digest and disrupt protein and lipid structures. Following the overnight digestion, magnetic beads resuspended in lysis/binding solution were added to the samples. Magnetic bead-bound DNA was transferred to subsequent wash steps by a magnetic particle processor. Total DNA was eluted and stored at -20 °C (or -80 °C) before quantification by qPCR.

### Optimization of digestion parameters

Based on the variability between protocols used in published studies, we identified protease digestion, detergent solubilization, sample volume, and incubation temperature and duration as potential factors that might impact ccf-DNA recovery (8, 22, 25, 31, 33–37). Therefore, we tested these variables for both ccf-mtDNA and ccf-nDNA (total ccf-DNA) recovery by testing concentrations, volume, and time to confirm and refine the effects on ccf-DNA yields (Fig. 2).

**Figure 2.**
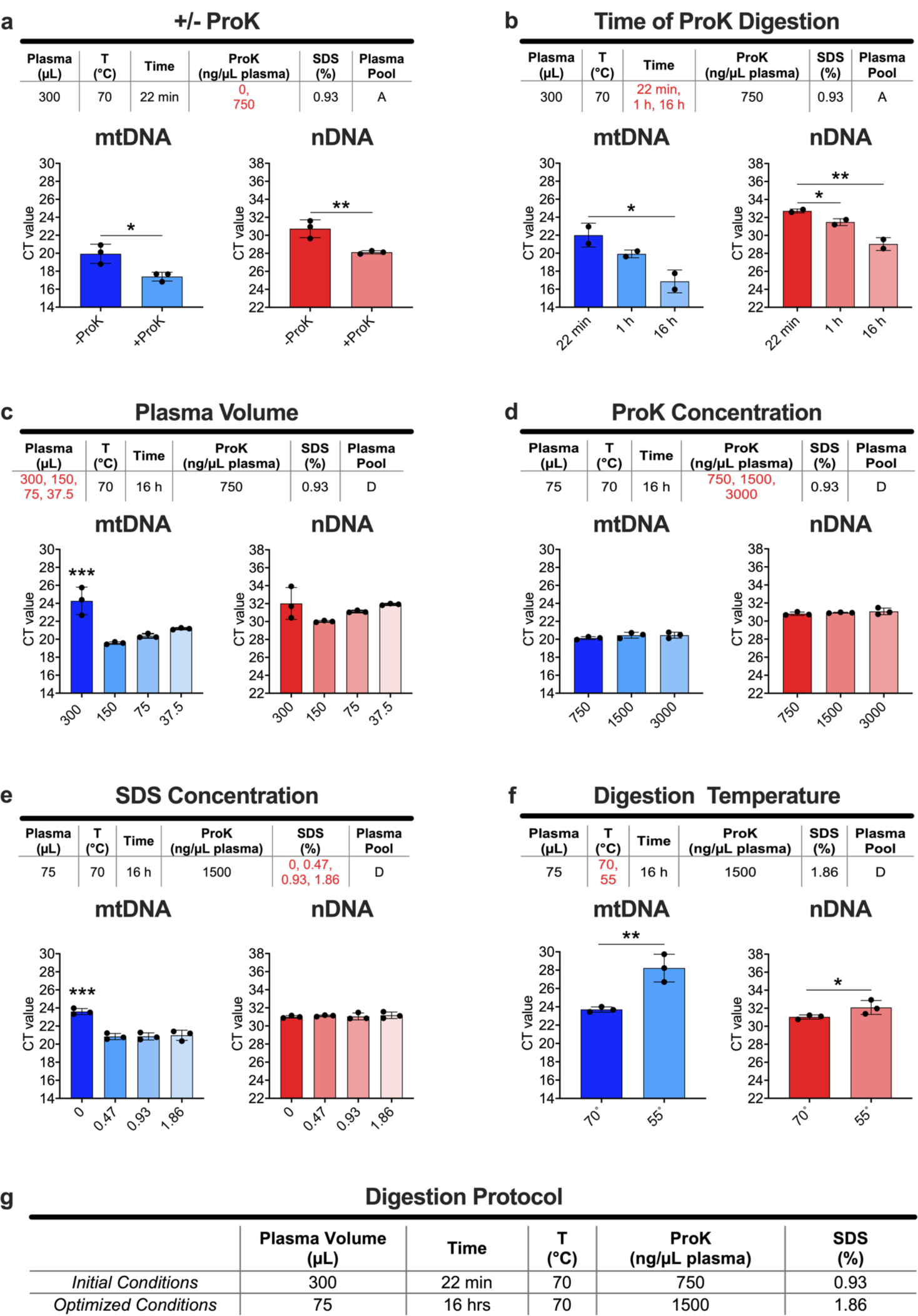
Optimization of digestion parameters. Depicted under each panel, the conditions tested include: (a) addition of ProK, (b) time of digestion, (c) plasma volume, (d) ProK concentration, (e) addition and final concentration of SDS, and (f) digestion temperature. The final conditions are summarized in (g). Variables color red are tested for contribution to DNA recovery. The plasma pool used for each experiment is indicated for each panel. Extractions were performed either by (a, b) the DynaMag-2 magnet or (c-f) Script 0 (at ambient temperature) in triplicate for each condition except for (b), which was done in duplicate. mtDNA and nDNA quantification are presented as mean CT values (log units) ± standard deviation. All reactions were run using a final volume of 8 μL with 3.2 μL of template DNA except (a) and (e), which were run in a final volume of 10 μL with 3.2 μL of template DNA. Statistical analysis was performed using (a, f) one-tailed, unpaired t-tests or (b-e) ordinary one-way ANOVA with Tukey’s multiple comparisons test between all conditions (p-values: * < 0.05, ** < 0.01, *** < 0.001, **** < 0.0001). In (c) and (e), all *post hoc* comparisons to the left-most condition are significant; however, for visual simplicity, only the p-value representing the ordinary one-way ANOVA is shown.

Plasma protein solubilization and digestion have been suggested to release ccf-DNA and increase recovery (25, 38). In initial experiments, we found that digestion with ProK for 22 min significantly increased total DNA recovery compared to digestions without ProK (Fig. 2a). Note that smaller CT values indicate greater DNA recovery. Time of digestion could also contribute to the recovery. Our previously described ethanol precipitation method for DNA isolation (9, 39, 40) digested tissue or plasma in ProK and SDS for 16 h rather than the 22 min recommended by the manufacturer. Hence, in an additional study, we compared the DNA yields from three time points: 22 min, 1 h, and 16 h (Fig. 2b). We found that recovery of mtDNA and nDNA improved progressively with longer incubation times (Fig. 2b).

It has been general practice to use large volumes of plasma (i.e. 0.5-2 mL) to isolate DNA, possibly to support downstream applications (25, 33, 41). However, no clear guidelines have been established with regards to the volume of plasma that should be digested, nor evidence of whether this parameter affects DNA recovery. We have not identified studies that isolated DNA from less than the manufacturer’s recommended volume. To address this, we tested DNA recovery from several plasma volumes (300, 150, 75, and 37.5 µL) (Fig. 2c). We wanted to determine the effect on ccf-DNA recovery when plasma volumes were decreased but extraction parameters were unchanged. ProK and SDS were added in the same proportion across the samples tested. However, the downstream extraction solution volumes were based on 300 µL of digested plasma (Script 0; Table 1), and thus, not adjusted for plasma volume. Surprisingly, reducing the plasma volume from 300 to 150 µL significantly increased mtDNA recovery (an 8-fold increase on average). Thereafter, mtDNA recovery was proportional to the input at all volumes ≤ 150 µL (Fig. 2c). Recovery of nDNA showed a similar trend (a 4-fold increase on average) but was not significant. To ensure that we were not close to the volume transition point that occurred between 300 and 150 µL, we chose 75 µL as our standard plasma volume. This has the added advantage of preserving valuable samples for additional studies.

**Table 1.** Summary of ccf-DNA extraction scripts used by this study. Each step in the process included solution volume(s) (µL), mix time (min), and mix speed variables. Beads were collected by the magnet and transferred to the next step as indicated. Temperature (24 °C) was an additional variable that was superimposed on Scripts 1 and 2, when applicable.

|  |  |  | Script |  |
| --- | --- | --- | --- | --- |
| Step | Process | 0 | 1 | 2 |
| 1 | Lysis/Binding Solution (μL) | 405 | 125 |  |
|  | Magnetic Beads (μL) | 10 | 3 | 5 |
|  | Mix time (min) [speed] | 6:00 [fast] | 6:00 [fast]* | 1:00 [fast]*<br>8:00 [medium]* |
|  | Number of bead collections | 2 |  | 1 |
| 2 | Wash 1 (μL) | 450 | 265 |  |
|  | Mix time (min) [speed] | 1:00 [fast] | 1:00 [fast]* | 1:00 [medium]* |
|  | Number of bead collections | 1 |  |  |
| 3 | Wash 2 (μL) | 900 | 475 |  |
|  | Mix time (min) [speed] | 1:00 [fast] | 1:00 [fast]* |  |
|  | Number of bead collections | 1 |  |  |
| 4 | Wash 3 (μL) | 200 |  |  |
|  | Mix time (min) [speed] | 00:30 [medium] | 00:30 [medium]* |  |
|  | Number of bead collections | 1 |  |  |
| 5 | Elution Solution (μL) | 60 |  |  |
|  | Mix time (min) [speed] | 6:00 [medium] | 6:00 [medium]* |  |
|  |  | Activate magnet to collect beads |  |  |
|  |  | 2:00 [slow] |  |  |

\* Indicates where uniform temperature (24°C) is applied to the extraction plates, when applicable

We next wanted to determine whether our current concentrations of ProK and SDS were resulting in maximal DNA yields. For ProK, we tested three concentrations (750, 1,500, and 3,000 ng ProK/µL plasma) (Fig. 2d) and found that increasing the concentration of ProK did not affect total ccf-DNA yields (Fig. 2d). In the above experiments, we had been using a final volume of 0.93% SDS in the plasma digestion. We tested whether this was an adequate or limiting amount by using four final concentrations of SDS: 0, 0.47, 0.93, and 1.86% (Fig. 2e). We found a significant increase in mtDNA recovery but no change to nDNA recovery when SDS was added, and that the increase in mtDNA was independent of the SDS concentration (Fig. 2e). To our knowledge, the SDS requirement for maximal ccf-mtDNA recovery has not previously been reported. Based on these findings, we chose 1,500 ng ProK/µL plasma and a final concentration of 1.86% SDS as our standard parameters, selected primarily to ensure adequate pipetting volume and limit variation in later experiments.

In addition to time, the temperature of ProK digestions are variable across different protocols, ranging from 37 °C to 60 °C (7, 8, 10, 22, 24, 25, 31, 33–37, 42, 43). Thus, we tested the impact of digestion temperature on plasma DNA recovery. We compared DNA yields obtained when plasma was incubated at the temperature recommended by the manufacturer (70 °C) and the temperature of our ethanol precipitation-based method (55 °C), which was within the range of temperatures that were previously published. When the digestion temperature was decreased from 70 °C to 55 °C, total ccf-DNA significantly decreased (Fig. 2f). At this point, our data suggest that incubating 75 µL of plasma with 1,500 ng ProK/µL plasma and 1.86% SDS for 16 h at 70 °C (Fig. 2g) results in increased yields.

### Optimization of extraction parameters

When identifying variables that could influence DNA yields, we selected extraction reagent volumes and MPP mixing parameters from our existing protocol as variables that could potentially limit DNA recovery. Because we found that a reduced plasma volume (75 µL as opposed to 300 µL) achieved better mtDNA recovery, we wanted to determine whether a proportional reduction in reagent volumes used during extraction would impact DNA yields. To begin, we extracted DNA from 75 µL of digested plasma using the original Script 0, whose extraction reagent volumes were based on 300 µL of digested plasma, and Script 1, whose volumes were based on 75 µL of plasma. Scripts 0 and 1 refer to the programmed actions of the MPP (i.e. the mixing and transferring of DNA-bound magnetic beads between plates) and the reagents that accompany each step. While all the actions were identical between Scripts 0 and 1, several Script 1 volumes were reduced to evaluate their cumulative effect on DNA recovery. We only modified the volumes of LBS+MB, Wash 1 (commercial solution), and Wash 2 (80% ethanol) (Table 1). We found that the volumes used in Script 1 did not negatively impact yields (Fig. 3a). In fact, mtDNA yields were unaffected while nDNA recovery improved.

**Figure 3.**
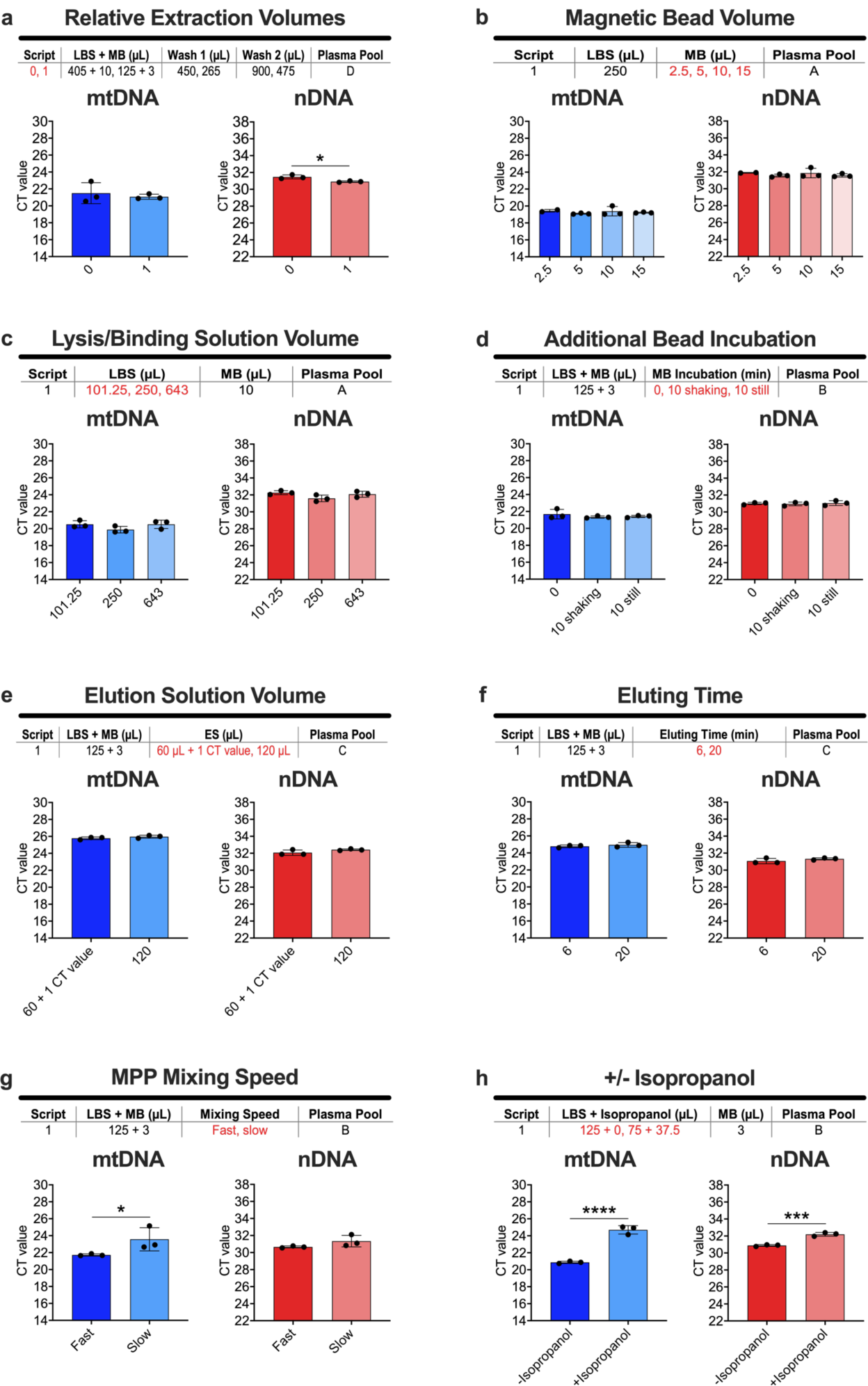
Optimization of extraction parameters. Depicted under each panel, the conditions tested include: (a) relative extraction volumes, (d) additional incubation with magnetic beads, (e) the addition of isopropanol to the lysis/binding and magnetic bead solution, and (f) MPP mixing speed, which were assessed for their impact on total DNA recovery. Varying volumes of (b) lysis/binding solution, (c) magnetic beads, and (g) elution solution, in addition to (h) eluting time, were evaluated to determine their points of saturation. Variables colored red are tested for their contribution to DNA recovery. The plasma pool used for each experiment is indicated. These extractions were performed using (a) Script 0 or (a-h) Script 1 (both at ambient temperature) in triplicate for each condition. mtDNA and nDNA quantification are presented as mean CT values (log units) ± standard deviation. All reactions were run using a final volume of 8 µL with 3.2 µL of template DNA except (b) and (c), which were run in a final volume of 10 µL with 1 µL of template DNA. Statistical analysis was performed using (a) a two-tailed, unpaired t-test, (e-h) one-tailed, unpaired t-tests, or (b-d) ordinary one-way ANOVA with Tukey’s multiple comparisons test between all conditions (p-values: * < 0.05, ** < 0.01, *** < 0.001, **** < 0.0001).

To confirm the results of Script 1, we tested MB or LBS volumes separately to evaluate their potential in limiting DNA yields. It has been previously demonstrated that MB were not easily saturated with DNA extracted from cells (32). However, MB saturation has not been investigated with regards to ccf-DNA isolations. As such, we extracted DNA from digested plasma using several volumes of MB (2.5, 5, 10, and 15 µL). Despite decreasing the volume of MB, the amount of total ccf-DNA recovered remained unchanged, suggesting the MB were in excess for the volume of plasma used (Fig. 3b). Likewise, we have not identified studies that have determined whether the volume of LBS influenced recovery. To test whether increasing LBS improved recovery, we extracted DNA from plasma using several volumes of LBS (101.25, 250, and 643 µL) but a fixed amount of MB. Our results reinforce the conclusion made from the script test in Fig. 3a by showing that DNA recovery was unchanged despite a decreased volume of LBS, suggesting that LBS was not a limiting reagent (Fig. 3c). To conserve reagents, we decided to proceed with the volumes of MB and LBS established in Script 1 (Table 1).

While the binding capacity of MB is independent of its volume (Fig. 3b), it is not known whether the process in which DNA binds to MB is time dependent. We tested this by incubating digested plasma with MB+LBS for 10 min before loading the samples onto the MPP for extraction. This incubation period more than doubled the amount of time that the DNA was allowed to interact with the beads (Script 1 already allotted for 6 min of mixing; Table 1). The extra incubation time was either on a shaking platform or a stationary rack (10 min shaking or 10 min still, respectively) and was compared to no additional time (indicated as “0” in Fig. 3d). As neither the increased time of DNA incubation with MB nor the agitation of the samples before MPP extraction had an effect on ccf-DNA recovery (Fig. 3d), we concluded that the standard incubation time was sufficient to reach DNA binding equilibrium.

Among the variables that we had identified from other studies, the volume in which DNA was eluted, which ranged from 30 to 200 µL, was not consistent and could potentially be a significant determinant of yields (8, 22, 25, 31, 33–37). To determine if all the DNA was being eluted from the MB using our current volume (60 µL), we eluted DNA in both 60 and 120 µL of ES. Because we doubled the elution volume, we would expect that the DNA would be half as concentrated when equal volumes of eluted DNA were used as the template for qPCR amplification. Assuming 100% extraction and quantification efficiencies, 1 CT value represents a 2-fold change in DNA concentration; therefore, we added 1 CT to the 60 µL result to compare the extraction recoveries (Fig. 3e). We also wanted to determine whether more time would release more DNA from the MB. To test this, we increased the eluting time from 6 min to 20 min (Fig. 3f). Increasing both elution volume and time had no impact on recovery and were not modified in later experiments.

In our review of the literature, we identified mixing speeds and the addition of isopropanol during the DNA binding step as remaining extraction parameters. We noted that a slow mixing speed was recommended for DNA isolation from serum in a technical note released by the manufacturer. In this plasma study, we found that a slow mixing speed significantly reduced mtDNA yields but had no measured effect on nDNA recovery (Fig. 3g). Furthermore, adding isopropanol to the LBS (for a final concentration of 50%) has been used to isolate cell-free DNA from sputum and urine (44) to promote DNA precipitation onto the MB (45). When using the Thermo Fisher chemistry, we found that isopropanol decreased recovery of mtDNA and nDNA significantly in plasma (Fig. 3h). In summary, these studies reinforce the parameter selections in Script 1, with the added benefit of decreasing reagent consumption.

### Identifying the source of variation in 96-well plate experiments

The first part of this study identified chemical parameters that influenced quantitative ccf-DNA recovery. Now, the second aim addresses issues affecting its reproducibility in experiments with large sample numbers. In an initial test, we isolated DNA from the same plasma pool for five consecutive days and analyzed mtDNA and nDNA yields on the same qPCR plate. We found that there was significant variation among batches (Fig. S1a), suggesting that additional variables were yet to be identified. From this result, we designed a set of experiments to quantify the relative variation from three potential sources: digestion, extraction, and quantification.

To identify the amount of variation from a single plate, DNA was isolated and quantified from 96 identical plasma samples (Fig. 4a). Surprisingly, both mtDNA and nDNA recovery were highly variable, with a standard deviation of 0.927 and 0.413, respectively (Fig. 4ai). To investigate whether this variation was due to well position, we plotted the difference in CT values relative to the median of the plate for each well (Fig. 4aii; raw data is presented in Fig. 1b-d). Importantly, while nDNA recovery was largely unaffected, less mtDNA was recovered around the edges and corners of the DWP (mtDNA edge effects). We did not know whether this variation arose from digestion, extraction, or qPCR quantification. We proceeded to work backwards from DNA quantification to identify the source of variation.

**Figure 4.**
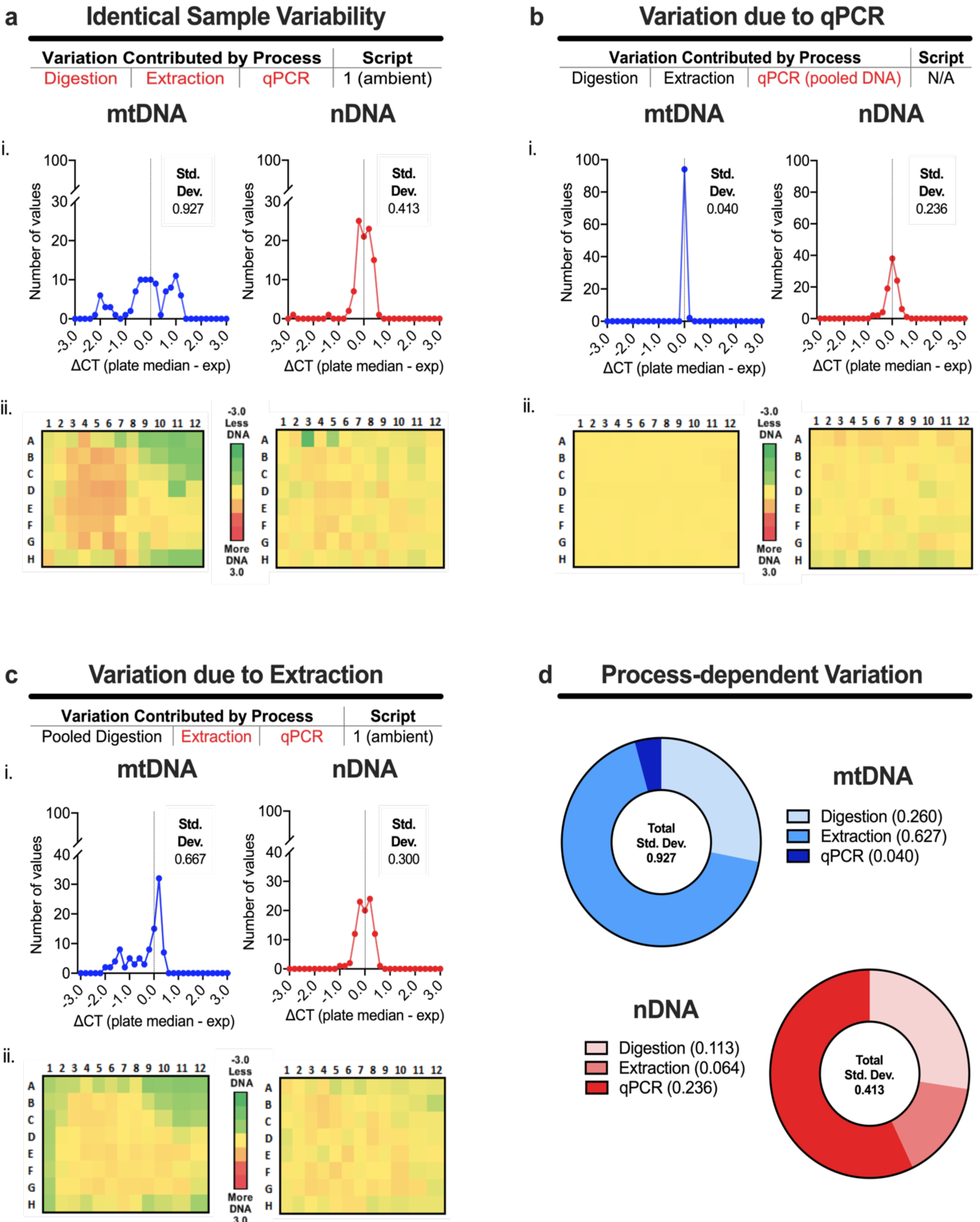
Identifying the source of variation during 96-well plate experiments. Depicted under the title of each panel are the steps in the process required to isolate and quantify ccf-DNA. When colored red, the step is being tested for its contribution to plate-wide variation across an identical sample isolated and/or quantified 96 times. To determine the combined variation contributed by plasma digestion, DNA extraction, and qPCR component steps, (a) ccf-DNA from plasma were digested overnight, extracted using Script 1 (at ambient temperature), and quantified by duplex qPCR (n = 96). To determine variation contributed by qPCR alone, (b) previously isolated ccf-DNA was pooled and quantified by duplex qPCR (n = 96). To determine the variation contributed by extraction alone, (c) pooled, digested plasma was isolated using Script 1 (at ambient temperature) and quantified by duplex qPCR (n = 96). (d) The pie charts summarize the calculated variation contributed by each process (based on the total standard deviation presented in (a)) on mtDNA and nDNA recovery. (a-c) ΔCT (plate median -experimental value [exp]) was calculated for each well and used in (i) to visualize the distribution of variation for 96 samples and in (ii) to determine the location of variation. In panel (ii), positional effects are visualized by assigning a color to ΔCT values ranging from -3.0 (less DNA, green) to 3.0 (more DNA, red), relative to 0.0 (plate median, yellow). Positional edge effects originating during extraction reduced mtDNA recovery only. qPCR reactions were run using a final volume of 8 µL with 3.2 µL of template DNA.

To determine how much of the variation across the plate was contributed by qPCR, we used pooled DNA from previous isolations and aliquoted it directly into a PCR plate to act as the source of template DNA. This ensured that the amount of DNA in all samples were identical. After qPCR quantification, mtDNA and nDNA variation were undetected, and the edge effects unique to mtDNA recovery were not present (Fig. 4b). We concluded that the variation did not arise from the qPCR assay itself or its associated liquid handling parameters (Fig. 4bii). We noted that nDNA CT values have a higher standard deviation than mtDNA, which would be expected because of the lower concentrations of template nDNA (46).

To determine the variation inherent to extraction, 96 samples of digested plasma were pooled, then aliquoted into a 96 DWP for extraction. This approach eliminated digestion as a variable. Subsequent qPCR analysis showed that for mtDNA recovery, extraction contributed about two-thirds of the total variation across the plate (Fig. 4ci). Variation in mtDNA recovery due to the digestion step was estimated by subtraction to be the remaining one-third of the total variation (Fig. 4d). Notably, edge effects in the distribution of mtDNA values clearly persisted when digestion variation was eliminated, thus isolating the extraction step as the primary source of mtDNA variation (Fig. 4cii). On the other hand, variation in nDNA recovery was modest (Fig. 4d). Because mtDNA recovery was highly variable due to positional effects, the results prompted further review of extraction parameters.

### Modifications to script parameters and plate temperature dramatically improved mtDNA recovery and reduced edge effects

The edge effects that occur during ccf-DNA extraction could arise from temperature gradients in or across the DNA extraction plates. To determine whether variability was affected by such gradients, DNA was extracted with the MPP’s internal heating element either turned off (an ambient temperature of 23 °C) or set to 24 °C for all steps. Variability in mtDNA recovery was significantly reduced when using warmed plates compared to ambient temperature (Fig. 5a), but some diminished mtDNA recovery remained localized to the top right corner of the DWP (Fig. 5bii; raw data is presented in Fig. S2). To determine if further elevating the temperature would result in even greater reductions in standard deviation and edge effects, DNA was extracted with the heating element set to 30 °C. However, the beneficial effects of warming on variability and positional effects were lost (Fig. S3) and higher temperatures were not pursued.

**Figure 5.**
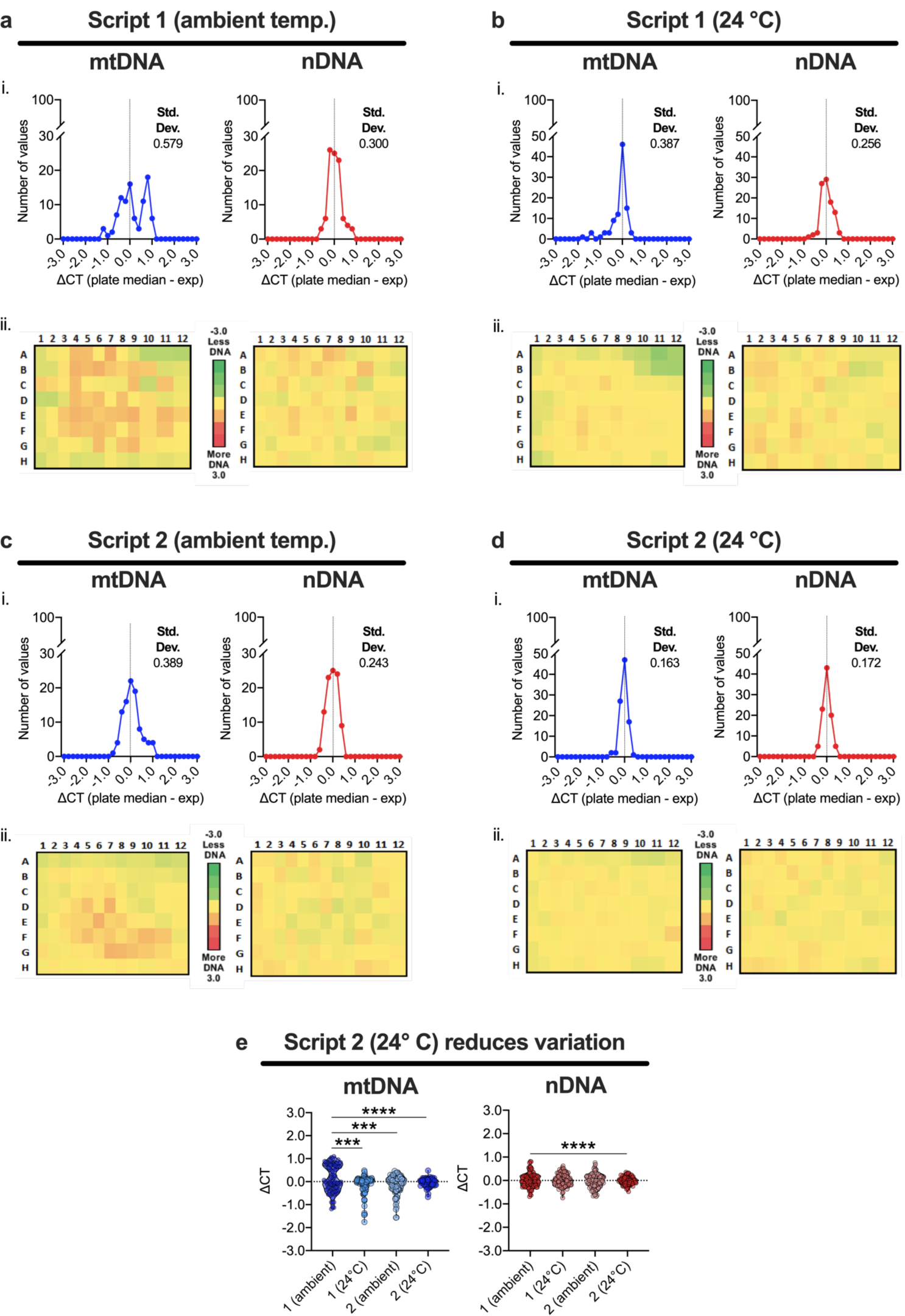
Modifications to script parameters and MPP temperature reduced the variation between wells. Each panel title (a-b, d-e) denotes the script being tested and the temperature (ambient or 24 °C) of extraction. Four plates of pooled, digested plasma were used to evaluate (a, c) the effects of a modified script and (b, d) uniform temperature (24 °C) on (e) the distribution of values across each plate. (a-d) ΔCT (plate median - experimental value [exp]) was calculated for each well and used in (i) to visualize the distribution of variation for 96 samples and in (ii) to determine the source of variation. In panel (ii), positional effects are visualized by assigning a color to ΔCT values ranging from -3.0 (less DNA, green) to 3.0 (more DNA, red), relative to 0.0 (plate median, yellow). mtDNA and nDNA quantification are presented as mean CT values (log units) ± standard deviation. (e) Violin plots were used to compare the distribution of variation of the four plates. qPCR reactions were run using a final volume of 8 µL with 3.2 µL of template DNA. Statistical analysis was performed using (e) F tests between 1 (ambient) and all other conditions (p-values: * < 0.05, ** < 0.01, *** < 0.001,

We were also concerned that MB processing events, such as a second collection of beads, were contributing to increased variation in the data. At this point, the manufacturer recommended modifications to several MPP script parameters (Script 2; Table 1), such as a single bead transfer between the digested plasma and Wash 1 plates and increased mixing times. When DNA was extracted using Script 2 at ambient temperature, mtDNA variation was improved relative to Script 1, but positional effects were still present (Fig. 5cii). We then combined the use of Script 2 and 24 °C MPP plate heating to determine if the improvements were cumulative. Remarkably, we found that when DNA was extracted and quantified from pooled, digested plasma using Script 2 at 24 °C that the mtDNA edge effects were visually undetectable (Fig. 5dii) and the variation among wells was significantly decreased for both mtDNA and nDNA (Fig. 5e). Thus, the data supported the implementation of Script 2 at 24 °C for future DNA extractions.

### Liquid handling characteristics that further improved mtDNA recovery

We next wanted to evaluate ccf-DNA yields when the entire process (plasma digestion, ccf-DNA extraction, and qPCR) was performed using complete automation. However, in preliminary experiments we noticed variable volumes of ProK and SDS remaining across wells of the 8-well PCR strips used by the LiHa to contain these reagents (Axygen Scientific, Union City, CA, USA). Liquid properties, such as viscosity, impact the accuracy in which reagents are dispensed by both manual pipettes and automated liquid handlers (47). Such liquids include SDS and glycerol, the main component of our resuspended ProK. Because ProK and SDS are crucial for effective plasma digestion (Fig. 2), we wanted to evaluate the reproducibility of the LiHa when dispensing these reagents. To achieve this, fluorescein (Thermo Fisher) diluted in ProK and SDS were separately dispensed by the LiHa. Fluorescence was measured using an Infinite F200 Fluorescence Microplate Reader (Tecan). Compared to fluorescein-spiked water, ProK and SDS dispenses varied significantly (Fig. S4). Further testing revealed that, when ProK and SDS were dispensed by the LiHa into plasma, mtDNA recovery was significantly reduced by 2 CT values, or four-fold less DNA, in four out of the eight plasma samples (Fig. 6a). Because nDNA recovery was unchanged (Fig. 6a), the data supports the visual confirmation (not pictured) that more SDS remained in four wells of the 8-well PCR strip (based on data from Fig. 2e). If ProK was not dispensed in adequate amounts, both mtDNA and nDNA recovery would have been significantly reduced (based on data from Fig. 2a).

**Figure 6.**
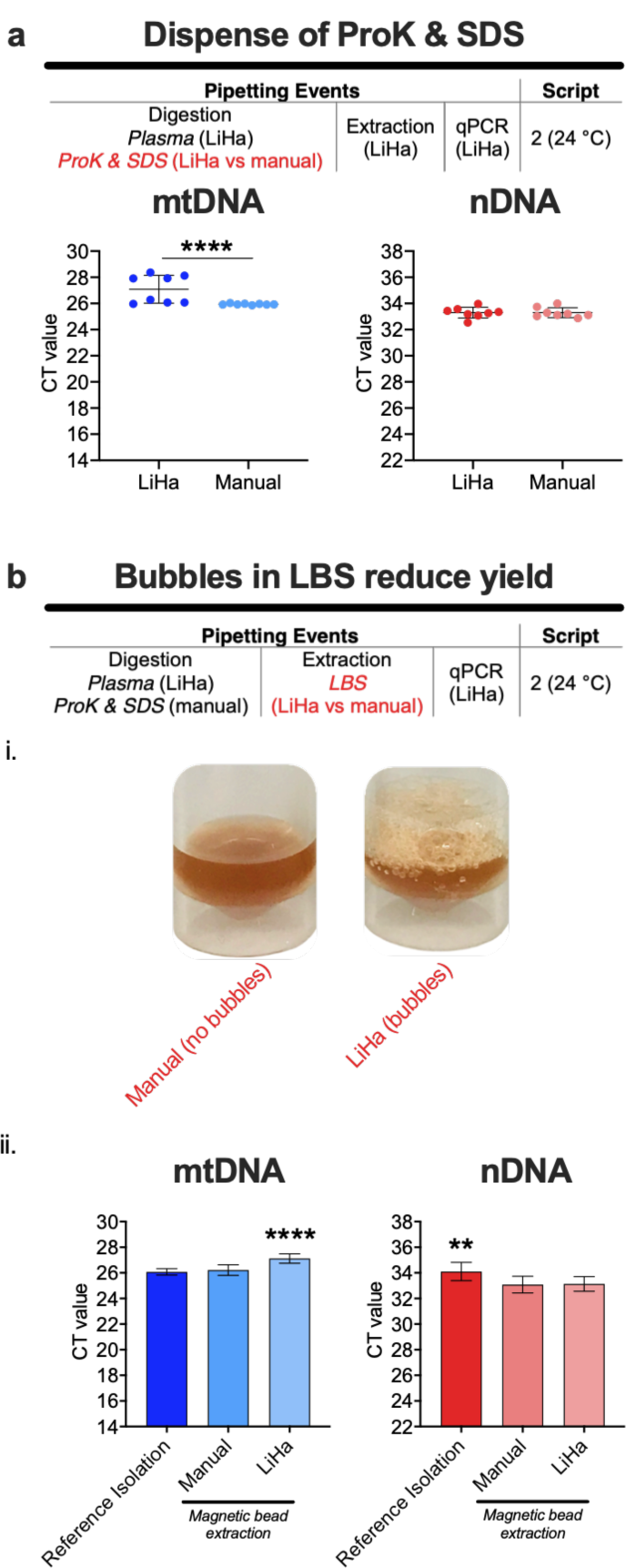
Liquid handling characteristics that further improved mtDNA recovery. Depicted under the title of each panel (a, b) are the steps in the process required to isolate and quantify ccf-DNA. When colored red, the step is being tested for its contribution to DNA isolation. To determine if the liquid handling settings were dispensing the digestion reagents accurately, (a) ccf-DNA was isolated and quantified by qPCR from two sets of plasma samples (n = 8) that differed in the ProK and SDS pipetting method (LiHa or manual). To determine if our liquid handling settings were pipetting the lysis/binding solution accurately, (b) ccf-DNA was isolated from two plates of plasma samples (n = 96) that differed in the pipetting method (LiHa or manual). In addition, ccf-DNA was isolated from plasma samples using our ethanol precipitation method as a reference benchmark for maximal ccf-DNA recovery (n = 6). Plasma pool D was used for all experiments. mtDNA and nDNA quantification are presented as mean CT values (log units) ± standard deviation. All reactions were run using a final volume of 8 µL with 3.2 µL of template DNA. Statistical analysis was performed using (a) an F test or (b) nonparametric Kruskal-Wallis test with Dunn’s multiple comparisons test between all conditions (p-values: * < 0.05, ** < 0.01, *** < 0.001, **** < 0.0001).

Other manufacturers’ methods have noted that bubbles in the LBS should be avoided as they reduce DNA yields (Omega Bio-tek, Inc., Norcross, GA, USA; Beckman Coulter Life Sciences, Indianapolis, IN, USA). Because we previously observed bubbles in LBS dispensed by the LiHa (Fig. 6bi), we compared the effects on yields when LBS was dispensed into digested plasma by the LiHa or a manual electronic pipette. While mtDNA recovery was similar to the ethanol precipitation-based reference isolation when a manual pipette was used, it was significantly decreased by the LiHa’s dispense (Fig. 6bii). However, nDNA yields were significantly better than those obtained by the reference isolation when either dispense method was used. Therefore, the data suggests that the manual dispense of SDS and LBS improved mtDNA recovery and should be incorporated into all subsequent plasma digestions and DNA extractions, respectively.

## Discussion

In this study, we focused on optimizing a commercially available MB-based isolation kit for the recovery of ccf-mtDNA and ccf-nDNA from plasma for incorporation into an automated platform. This study has unexpectedly uncovered flaws in the reproducibility of ccf-mtDNA isolation. Because the majority of research has focused on ccf-nDNA (more generally termed ccf-DNA), the manufacturer would not be aware of this limitation. Our intention was not to discredit the existing protocol (which worked well for ccf-nDNA), but to ensure that the assay served our purposes as a quantitative, high throughput method for both ccf-nDNA and ccf-mtDNA isolation. With ccf-mtDNA under investigation for its use as a clinical biomarker, additional studies will be necessary to find the optimal procedure. We encourage labs interested in achieving maximal ccf-mtDNA recovery to consider the results presented here for implementation into their own assays.

It has been widely accepted that plasma, rather than serum, should be used to evaluate ccf-DNA content. Studies have demonstrated that, during coagulation, white blood cell lysis increases the levels of nDNA in serum. Moreover, platelet activation during coagulation can also release mitochondria and mtDNA (48), and quantification of ccf-mtDNA in a parallel comparison of serum and plasma samples from the same individual is consistent with this claim (49). Therefore, immediate processing of whole blood to obtain plasma is preferable for isolating ccf-DNA, as both nDNA and mtDNA contamination are significantly reduced (27, 34, 49, 50). In our study, we heeded this recommendation and optimized our protocol using plasma. However, the lots used in this study had unique levels of total ccf-DNA content due to their various origins. Thus, when results are presented as mean CT values, the plasma pool used for each experiment is indicated under the panel title to prevent comparison between results obtained using different plasma pools.

The role that plasma digestion plays in quantitative ccf-DNA isolation is largely unclear and, as a result, protocols vary in their recommendations concerning plasma volume, temperature, and duration (7, 8, 10, 22, 24, 25, 31, 33–37, 42, 43), with some protocols disregarding both plasma digestion (25) and DNA extraction entirely (21). However, one study hypothesized that proteolytic digestion increases isolation efficiencies by releasing ccf-DNA that is bound to proteins or contained in extracellular vesicles (25). Our data, like that of Xue et al. (2009), suggested that the absence of ProK significantly decreased total ccf-DNA yields. Likewise, mtDNA recovery significantly decreased when SDS was not present, thus supporting studies that have demonstrated that some mtDNA is contained within exosomes in the circulation (51, 52). Further studies are required to determine how the DNAs isolated in these contexts differ in source and characteristic.

It has been assumed that volumes of plasma ≥ 500 µL are required to recover adequate DNA yields for downstream applications (33). However, we demonstrated that the manufacturer’s recommended plasma volume (300 µL) does not produce optimal mtDNA yields in this automated workflow. When we isolated DNA from plasma volumes ≤ 150 µL, we observed improved mtDNA recovery for reasons that are not clear. In our previously described ethanol precipitation method for ccf-DNA isolation (9, 39, 40), plasma was digested at 55 °C, an optimal temperature for that chemistry. In the current MB chemistry, total DNA yields were improved at 70 °C relative to 55 °C with the same ProK, suggesting that digestion temperatures need to be optimized for each specific chemistry used. While also required to achieve maximal recovery, one downfall of the optimized digestion conditions was the increase in digestion time (16 h), and therefore, an overall lengthened isolation process. With breaking the process over two days, we find that a similar amount of work can be completed by rearranging the schedule. The improved conditions have proved useful for our purposes as a method for obtaining maximal ccf-DNA recovery for qPCR assessment. However, that does not mean that for other applications, such as detecting the presence of specific DNA sequences or sequencing-based methods, that these specific digestion parameters would be sufficient. Thus, this preparation should be carefully tested for any other downstream applications.

Based on the review of the literature, we expected that improved DNA recovery would come from chemistry-specific extraction parameters. However, we were surprised to find that mechanical aspects made sizeable differences in the performance of the protocol when assessed for its reproducibility during high throughput experiments, rather than the reagents themselves. For example, mtDNA but not nDNA recovery was subject to edge effects, resulting in increased isolation variability and decreased yields when ambient temperatures were not controlled. The characteristics of mtDNA that contribute to this temperature effect are not clear. Furthermore, we thought that modifications to the MPP script, such as decreasing the number of bead transfers and mixing characteristics to create Script 2 (Table 1), would have no effect on mtDNA edge effects and possibly decrease total DNA yield. Instead, it was somewhat helpful in edge effects and yield, and in combination with temperature control, eliminated the plate variation. We are unaware of any published descriptions of these effects for mtDNA isolation, either cellular or cell-free.

During the optimization of digestion parameters, we determined that ProK and SDS were required for quantitative DNA recovery, but their pipetting was not uniform when using the LiHa. Recognizing that further liquid class optimization is required, we had already taken several steps to prevent this confounder. First, our solutions were centrifuged to remove bubbles. Second, we performed deep pipetting to reduce the likelihood of air aspiration. Third, we greatly decreased the aspiration speed to 5 µL/sec for both SDS and ProK. Lastly, we knew that pipetting of SDS and ProK was tolerant of at least 50% pipetting error as we had chosen larger than minimal volumes for the final protocol. The digital repeater is a reasonable alternative to automated pipetting in this situation, as liquid dispensing can be validated visually and does not significantly increase pipetting time. Furthermore, while we anticipated some variability in liquid handling of SDS, the generation of bubbles in the LBS solution and its reduction of mtDNA yields was an unforeseen issue that also was easily mitigated by manual pipetting. Regardless, we want to highlight that the optimization of liquid handling settings for critical reagents, such as SDS and LBS, should be taken into consideration for labs that desire full automation.

While some studies have expressed concern about the ease of implementing an automated workstation into their labs, including an extensive learning curve and empirical optimization (25), we recognize that many institutions may not have the funds or need for an automated methodology (32). Fortunately, the chemistry surrounding our protocol is compatible with microfuge tube magnets (such as the DynaMag-2 magnet used in some of these studies), which bypasses automated MPPs, requires minimal training, and is appropriate for small-scale experimentation. However, for implementing ccf-DNA measurements in routine clinical practice or other high throughput applications, automation would limit human error and reduce processing time. As the study of ccf-DNA abundance expands to other applications, reliable process automation is an important tool to enable the identification of associated clinical parameters.

When using a MB-based ccf-DNA isolation approach, there are overlapping parameters of plasma digestion that should be applied to achieve maximal mtDNA and nDNA recovery. These include the addition of ProK and an extended digestion period of 16 h at 70 °C. However, incubation with SDS only improved mtDNA yields. We have also demonstrated that edge effects originating during extraction significantly reduced mtDNA recovery with little to no effect on nDNA isolation. Application of uniform temperature (24 °C) across all extraction plates coupled with an improved script reduced edge effects and improved total ccf-DNA variation between wells. Taken together, these improvements resulted in equal or greater yields compared to the common ethanol precipitation-based method. With ccf-mtDNA on the rise as a potential clinical biomarker, the need to establish a uniform methodology for its isolation is pressing (53). To our knowledge, this is the first high-throughput method optimized for ccf-mtDNA and ccf-nDNA recovery. Because we have identified issues in the recovery and reproducibility of an automated, MB-based ccf-DNA isolation protocol and outlined initial solutions, this study serves as an important starting point for the implementation of ccf-mtDNA and ccf-nDNA analysis in clinical studies.

## Experimental procedures

### Plasma samples

Five distinct plasma pools (A-E) were used for this study. To obtain plasma pool A, venous blood was collected using EDTA-containing vacutainer tubes (Cat. # 367835; Becton, Dickinson and Company, Franklin Lakes, NJ, USA) and centrifuged immediately at 950 x *g* for 15 minutes (min) at 4 °C. Plasma was collected, pooled, aliquoted, and stored at -80 °C. Pools B-E were purchased (BioIVT, Hicksville, NY, USA). Each lot was aliquoted and stored at -80 °C.

### Liquid handling system description

The workstation used in this study was the Freedom EVO 150 automated liquid handler (Tecan, Männedorf, Switzerland). This workstation was equipped with an 8-channel fixed tip, water-based pipetting system (LiHa) and a robotic arm (RoMa) controlled through the EVOware Standard software (EVOware; Ver. 2.3, Tecan). As appropriate, the tips were rinsed twice between pipetting events using 10% bleach and flushed at both washing stations to minimize sample cross-contamination. A KingFisher Presto (Thermo Fisher, Waltham, MA, USA) <u>m</u>agnetic <u>p</u>article <u>p</u>rocessor (MPP) was integrated adjacent to the workstation. To allow for complete automation, the RoMa was programmed to transport plates between the liquid handling platform and MPP turntable.

### Automated ccf-DNA isolation

#### Plasma digestion

Plasma, <u>Pro</u>teinase <u>K</u> (ProK; Thermo Fisher), and <u>s</u>odium <u>d</u>odecyl <u>s</u>ulfate (SDS; Boston BioProducts, Ashland, MA, USA) were dispensed into gasketed, screw-cap tubes (Sarstedt Inc., Nümbrecht, Germany) or a 96 <u>d</u>eep-<u>w</u>ell <u>p</u>late (DWP; Thermo Fisher) that was sealed with an adhesive PCR seal (Thermo Fisher) and covered with generic packaging tape. Samples were incubated in an Innova 44 Incubator Shaker (New Brunswick Scientific, Edison, NJ, USA) or on a digital orbital mixing chilling/heating dry bath (Torrey Pines Scientific, Carlsbad, CA, USA). Samples digested in tubes were manually transferred to a DWP for extraction. For some experiments, samples were pooled to eliminate variation due to digestion.

### Extraction of ccf-DNA from digested plasma

All samples were extracted using the MPP except for two initial studies that were extracted using the DynaMag-2 (Thermo Fisher) benchtop magnet (indicated in the figure legend). The reagents described below were loaded by the LiHa or a HandyStep digital repeater pipette (BrandTech Scientific, Inc., Essex, CT, USA) as indicated in the results. The MPP scripts (Table 1) were created in the BindIt 4.0.0.45 for KingFisher Instruments software (Thermo Fisher) and an automation interface to Tecan’s EVOware software executed each step of the script as programmed.

The extraction process used five different buffer incubations. To begin the extraction process, the digested plasma was removed from the incubator and left at room temperature (RT) for ∼15 min while the extraction consumables were placed on the platform. The digested plasma was centrifuged at 500 x *g* for 1 min before removing the seal and being placed on the platform. The user then launched the EVOware software and executed the extraction script. Key technical aspects of the extraction process, including reagent volumes and mixing characteristics, are summarized in Table 1. A DWP supporting a KingFisher 96 tip <u>c</u>omb (TC) for deep well magnets (Thermo Fisher) was loaded onto the MPP by the RoMa. The TC was loaded onto the magnetic head of the MPP and then the DWP was removed from the MPP platform by the RoMA. For most experiments, the LiHa dispensed Dynabeads MyOne Silane <u>m</u>agnetic <u>b</u>eads (MB; Thermo Fisher) re-suspended in MagMAX Cell Free DNA <u>L</u>ysis/<u>B</u>inding <u>S</u>olution (LBS; Thermo Fisher) into the DWP that contained the digested plasma. The RoMa loaded the plate containing digested plasma and MB+LBS onto the MPP. The samples were then mixed by the TC while the LiHa dispensed MagMAX Cell Free DNA Wash Solution (Wash 1; Thermo Fisher). After the RoMa loaded Wash 1 onto the MPP, the magnet collected the MB and bound DNA. The MPP transferred the beads to Wash 1 for subsequent mixing by the TC. The RoMa removed the DWP containing the residual digested plasma and the process of plate addition, mixing, and removal repeated for Washes 2, 3 (80% ethanol) and the MagMAX Cell Free DNA <u>E</u>lution <u>S</u>olution (ES; Thermo Fisher). All extraction reagents were dispensed into DWPs except ES, which was dispensed into a KingFisher 200 µL 96 well plate (Thermo Fisher). To release the DNA from the MB, the MB and ES were mixed by the TC and then the MB were captured by the magnet. The RoMA collected the ES plate (containing DNA) and placed it onto a magnetic plate (Alpaqua, Beverly, MA, USA) to capture any MB carryover. The ES solution was then transferred to a 96-well PCR Plate (Thermo Fisher) for storage and ended the automation. The plate was manually sealed with an adhesive PCR seal and stored at -20 °C.

### Ethanol precipitation-based reference isolation

For comparison, we isolated ccf-DNA using a previously described ProK digestion and ethanol precipitation method (9, 39, 40) with the minor modification of using ES for DNA resuspension. In brief, 75 µL of plasma were incubated at 55 °C for 16 h with 0.83 µL of 20% SDS, 7.58 µL of 5 M Tris-HCl (pH 8.5; Thermo Fisher), 0.76 µL of 5 M EDTA (Thermo Fisher), 3.12 µL of 5 M NaCl (Thermo Fisher), 2.62 µL of 20 mg/mL ProK, and 0.70 µL of 1% BME. After the overnight digestion, the samples were allowed to cool at RT for 5 min. The samples were then diluted 4.5:1 with pre-mixed digestion buffer containing 0.2% SDS, 100 mM Tris-HCl, 5 mM EDTA, 200 mM NaCl based on the concentration of the stock solutions described above (409.4 µL of digestion buffer for 90.6 µL of digested plasma). The digestion buffer was followed by 170 µL of 5 M NaCl and a five-minute incubation on ice to precipitate proteins. The samples were centrifuged at 26,000 x *g* for 15 min at 4 °C. The supernatant was transferred to a fresh tube to which 800 µL of cold 100% ethanol (Decon Labs Inc.) and 1 µL of GlycoBlue nucleic acid co-precipitant (15 mg/mL; Thermo Fisher) were added. The tubes were centrifuged again at 26,000 x *g* for 15 min at 4 °C. The supernatant was removed by aspiration and 500 µL of cold 70% ethanol was added. The tubes were centrifuged at 28,500 x *g* for 5 min at 4 °C. The supernatant was removed by aspiration and the pellet air dried at 37 °C for 15 min. 60 µL of ES was added to the tubes and incubated at 55 °C for 1 h. The resuspended DNA was stored at -20 °C.

### Quantitative polymerase chain reaction (qPCR)

ccf-mtDNA and ccf-nDNA abundance were measured simultaneously by duplex qPCR using TaqMan-based assays. These duplex reactions were run in triplicate (3 wells) on a QuantStudio 5 Real-Time PCR System (Thermo Fisher). One of two duplex assay pairings were used for each experiment; both of which have been previously validated (9, 54). The first pairing targeted mitochondrial-encoded human NADH:ubiquinone oxidoreductase core subunit 1 (ND1) and nuclear-encoded human beta-2-microglobulin (B2M). The second pairing targeted mitochondrial-encoded human NADH: ubiquinone oxidoreductase core subunit 4 (ND4) and nuclear-encoded human peptidylprolyl isomerase A (PPIA). Each target’s assay, composed of two primers and a fluorescent probe, were assembled as a 20X working solution according to the manufacturer’s protocol (Integrated DNA Technologies, Newark, New Jersey, USA) and stored at -20 °C.

For a duplex reaction, primer/probe ratios of 1:1 nmole (i.e. primer-limited) for mtDNA assays and 3:1 nmoles for nDNA assays were used. Thermocycling conditions were as follows: an initial denaturation step at 95 °C for 20 sec followed by 40 cycles of 95 °C for 1 sec, 63 °C [ND1/B2M] or 59 °C [ND4/PPIA] for 20 sec, and 60 °C for 20 sec. Assay sequences are in Table S1. Either 2X Luna Universal qPCR Master Mix (New England Biolabs, Ipswisch, ME, USA) or 2X TaqMan Fast Advance Master Mix (Thermo Fisher Scientific) were used in this study. The qPCR master mix and TaqMan assays for both mtDNA and nDNA targets were diluted to 1X in the final reaction volume. Reactions with a final volume of 8 µL contained 3.2 µL of template DNA; reactions with a final volume of 10 µL contained either 3.2 µL or 1 µL of template DNA. While the qPCR reaction size did not impact the results, the volumes are indicated in the figure legends.

### Statistical analysis

All statistical analyses were performed using GraphPad Prism (GraphPad Software, Ver. 8.3). Normality of distribution was tested in experiments with 96 biological replicates using the Anderson-Darling, D’Agostino & Pearson, Shapiro-Wilk, and Kolmogorov-Smirnov tests; nonparametric statistics were applied unless all tests confirmed normality, for which parametric tests were used. The statistical test used for each experiment is indicated in the figure legends. P-values < 0.05 were considered statistically significant.

## Data availability

All data are contained within the manuscript.

## Acknowledgements

The authors want to thank Xingwang Fang, PhD for his helpful discussions, Claudia Lagranha, PhD for reading the manuscript, and Kevin Redding for his input on figure design. Fig. 1, Fig. 1S, and the graphical abstract were created using BioRender.

## Abbreviations

ccf: circulating, cell-free;
DWP: deep well plate;
ES: Elution Solution;
exp: experimental value;
h: hours;
LBS: Lysis/Binding Solution;
LiHa: liquid handler;
MPP: magnetic particle processor;
min: minutes;
mtDNA: mitochondrial DNA;
nDNA: nuclear DNA
ProK: Proteinase K;
RoMa: robotic arm;
RT: room temperature;
SDS: sodium dodecyl sulfate;
sec: seconds;
TC: tip comb;
qPCR: quantitative polymerase chain reaction.

## Supporting Information

**Table S1.**
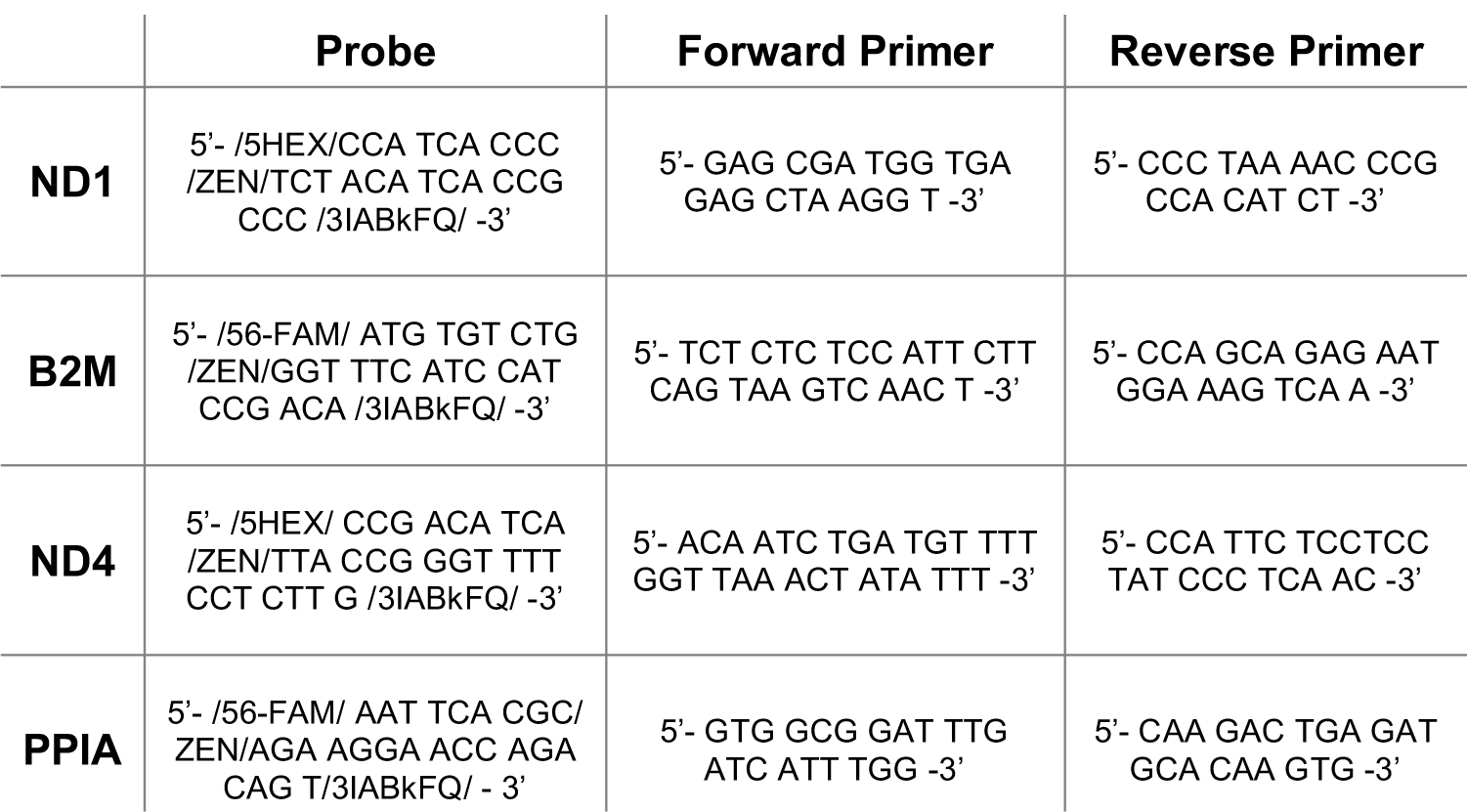
qPCR assay sequences. Two sets of duplex qPCR assays were used in this study. The first targeted ND1 and B2M while the second targeted ND4 and PPIA.

**Figure. S1.**
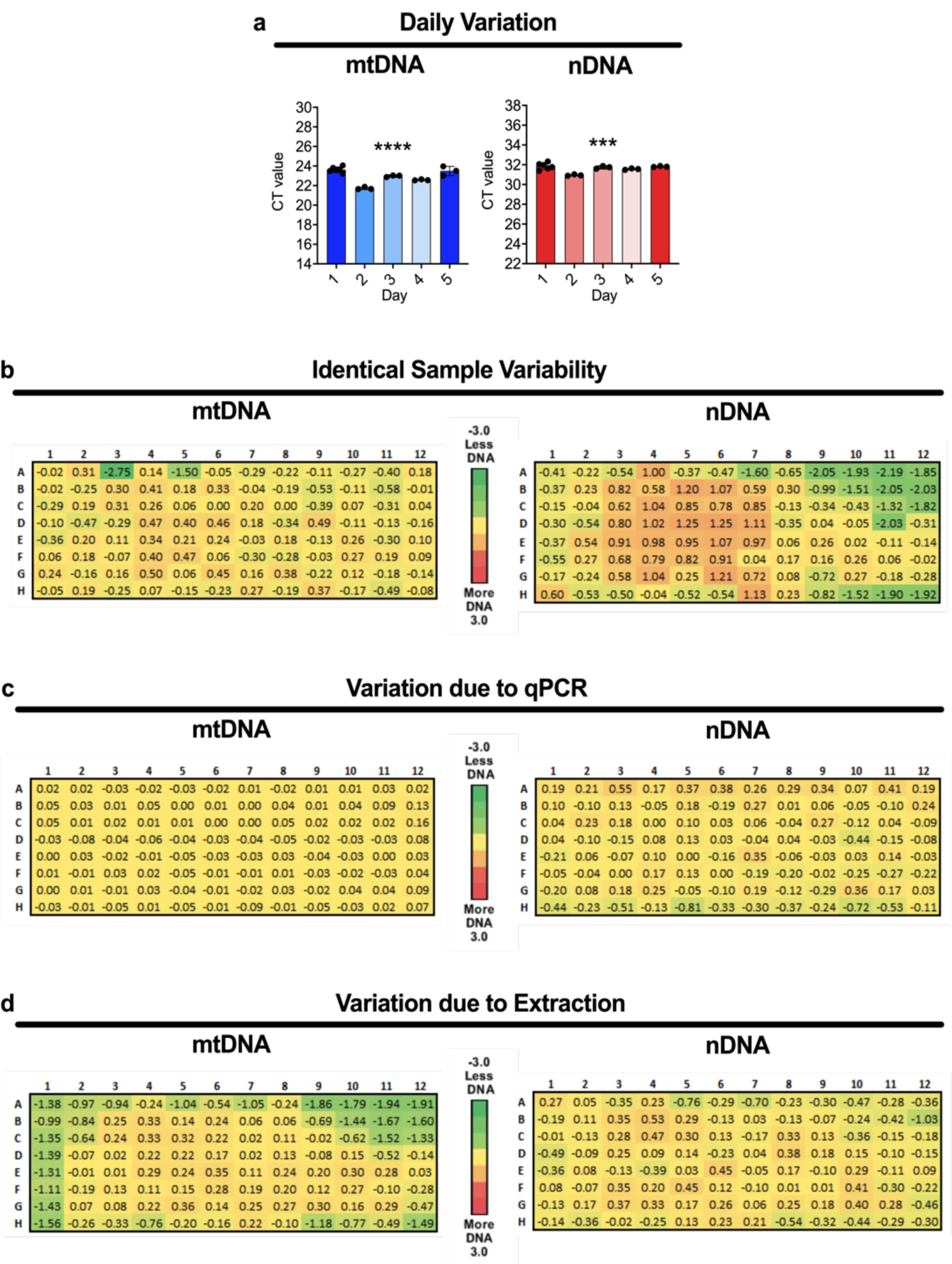
ccf-DNA variation. When several days of ccf-DNA isolations were quantified at once by duplex qPCR, (a) both mtDNA and nDNA yields were significantly variable between days. Statistical analysis was performed using ordinary one-way ANOVA (p-values: * < 0.05, ** < 0.01, *** < 0.001, **** < 0.0001). (b-d) Raw data corresponding to Fig. 5a-cii with mtDNA and nDNA presented as ΔCT (plate median – experimental values).

**Figure S2.**
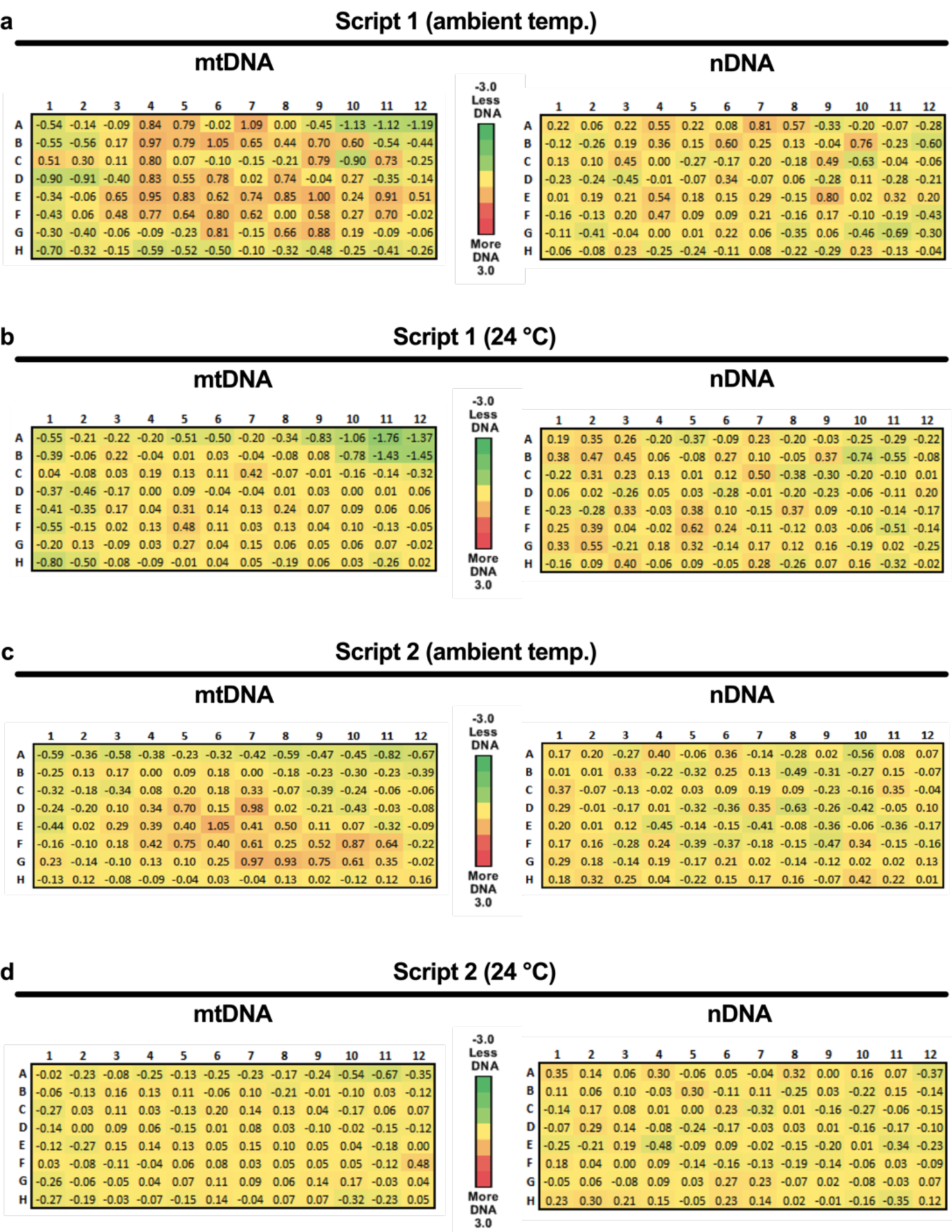
ΔCT (plate median – experimental values) have been added to the plate depictions corresponding to those in Fig. 6a-dii.

**Figure S3.**
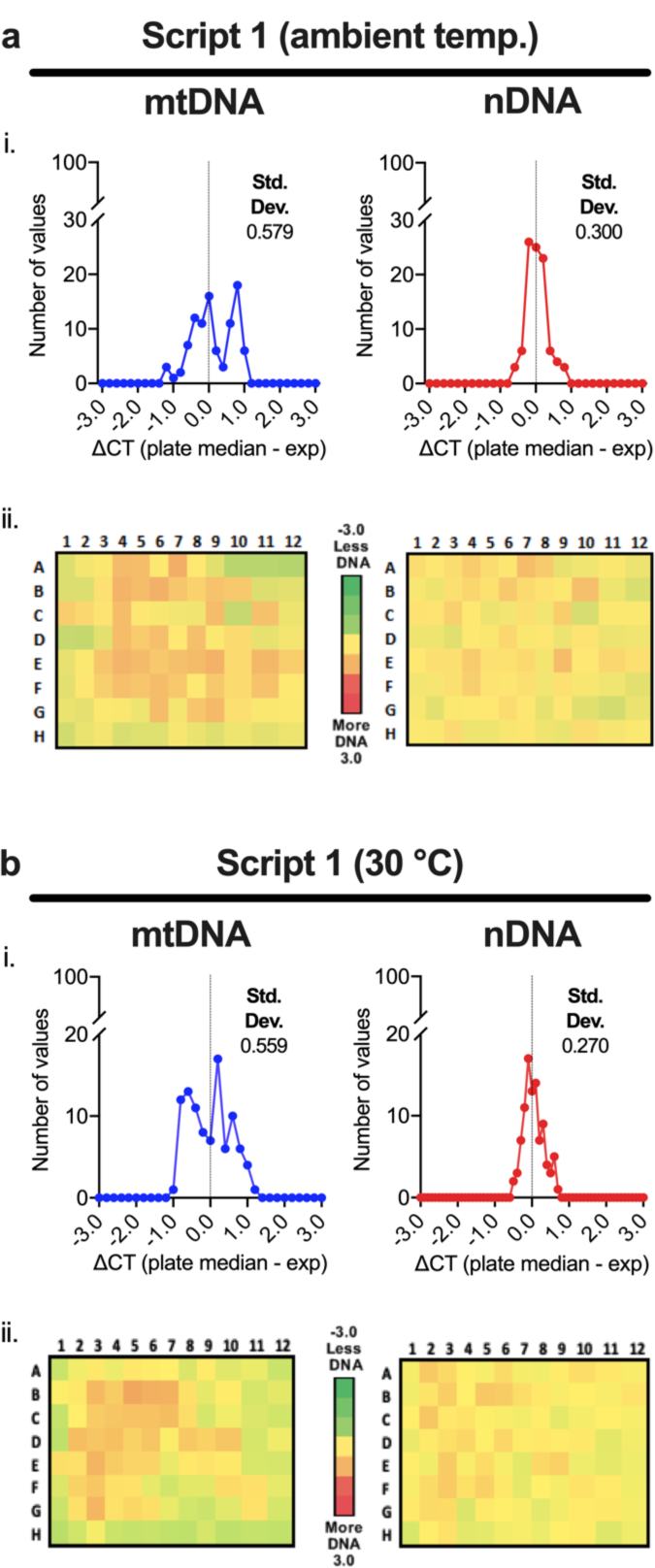
Applying 30 °C across all extraction plates did not improve standard deviation or edge effects. For this experiment, the temperature setting of the MPP was either off or at 30°C during the extraction process.

**Figure S4.**
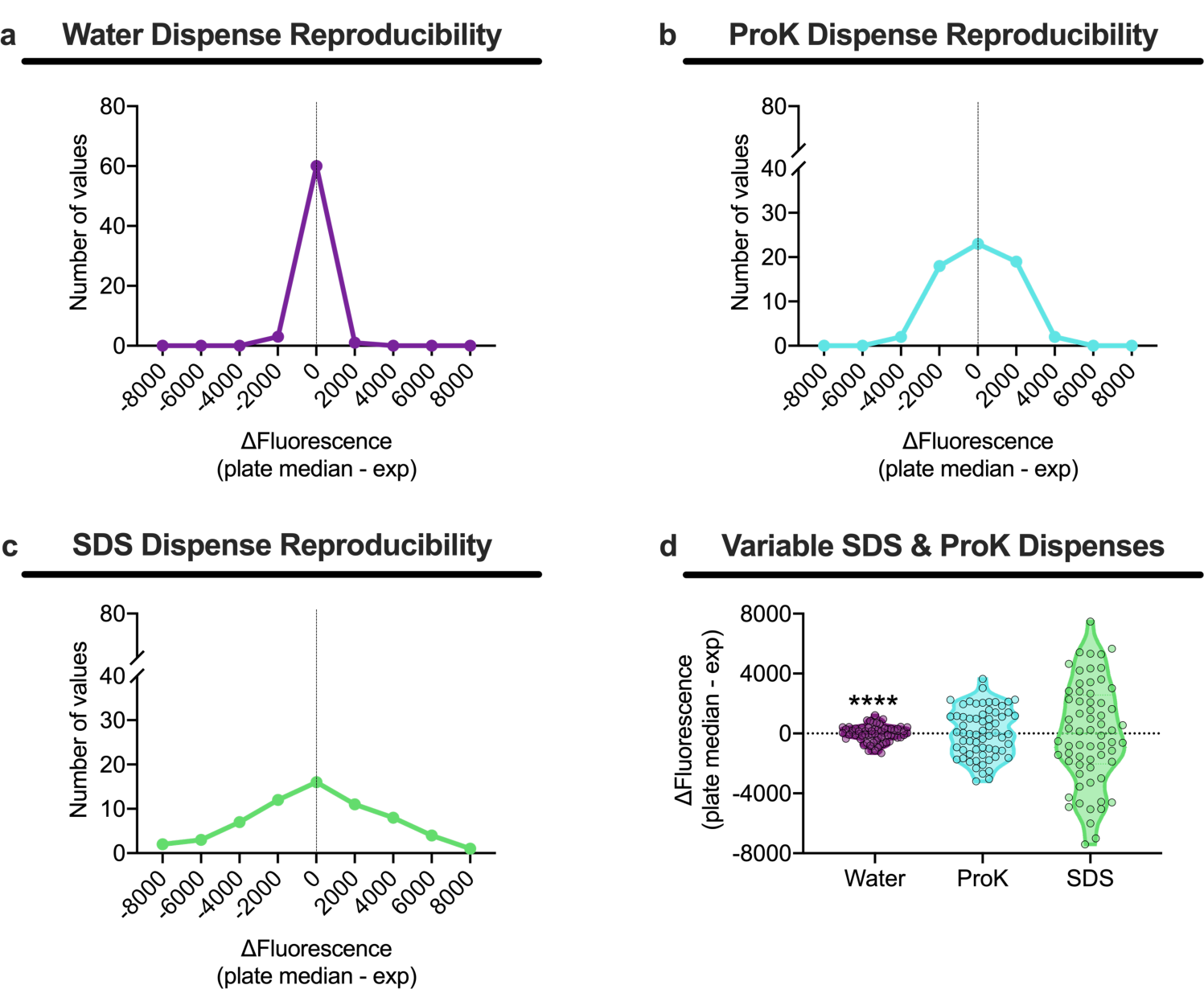
Volumes of ProK and SDS dispensed by the LiHa were variable. Fluorescein diluted in (a) water, (b) ProK, and (c) SDS was dispensed by the LiHa and fluorescence was measured (n = 64 per group). ΔFluorescence (plate median – exp) was calculated for each dispense. (d) Violin plots were used to compare the distribution of variation of the three conditions and statistical analysis was performed using multiple F tests where water was used as the reference condition (p-values: * < 0.05, ** < 0.01, *** < 0.001, **** < 0.0001).

## References

1. Zhong, S., Ng, M. C., Lo, Y. M., Chan, J. C., and Johnson, P. J. (2000) Presence of mitochondrial tRNA(Leu(UUR)) A to G 3243 mutation in DNA extracted from serum and plasma of patients with type 2 diabetes mellitus. J. Clin. Pathol. 53, 466–469

2. Nakahira, K., Kyung, S.-Y., Rogers, A. J., Gazourian, L., Youn, S., Massaro, A. F., Quintana, C., Osorio, J. C., Wang, Z., Zhao, Y., Lawler, L. A., Christie, J. D., Meyer, N. J., Mc Causland, F. R., Waikar, S. S., Waxman, A. B., Chung, R. T., Bueno, R., Rosas, I. O., Fredenburgh, L. E., Baron, R. M., Christiani, D. C., Hunninghake, G. M., and Choi, A. M. K. (2013) Circulating mitochondrial DNA in patients in the ICU as a marker of mortality: derivation and validation. PLoS Med. 10, e1001577. discussion e1001577

3. Gögenur, M., Burcharth, J., and Gögenur, I. (2017) The role of total cell-free DNA in predicting outcomes among trauma patients in the intensive care unit: a systematic review. Crit. Care 21, 14

4. Itagaki, K., Kaczmarek, E., Lee, Y. T., Tang, I. T., Isal, B., Adibnia, Y., Sandler, N., Grimm, M. J., Segal, B. H., Otterbein, L. E., and Hauser, C. J. (2015) Mitochondrial DNA released by trauma induces neutrophil extracellular traps. PLoS One 10, e0120549

5. Bhagirath, V. C., Dwivedi, D. J., and Liaw, P. C. (2015) Comparison of the proinflammatory and procoagulant properties of nuclear, mitochondrial, and bacterial DNA. Shock 44, 265–271

6. Zhang, Q., Raoof, M., Chen, Y., Sumi, Y., Sursal, T., Junger, W., Brohi, K., Itagaki, K., and Hauser, C. J. (2010) Circulating mitochondrial DAMPs cause inflammatory responses to injury. Nature 464, 104–107

7. Arshad, O., Gadawska, I., Sattha, B., Côté, H. C. F., Hsieh, A. Y. Y., and Canadian Institutes of Health Research Team on Cellular Aging and HIV Comorbidities in Women and Children (CARMA) (2018) Elevated Cell-Free Mitochondrial DNA in Filtered Plasma Is Associated With HIV Infection and Inflammation. J. Acquir. Immune Defic. Syndr. 78, 111–118

8. Lindqvist, D., Wolkowitz, O. M., Picard, M., Ohlsson, L., Bersani, F. S., Fernström, J., Westrin, Å., Hough, C. M., Lin, J., Reus, V. I., Epel, E. S., and Mellon, S. H. (2018) Circulating cell-free mitochondrial DNA, but not leukocyte mitochondrial DNA copy number, is elevated in major depressive disorder. Neuropsychopharmacology 43, 1557–1564

9. Trumpff, C., Marsland, A. L., Basualto-Alarcón, C., Martin, J. L., Carroll, J. E., Sturm, G., Vincent, A. E., Mosharov, E. V., Gu, Z., Kaufman, B. A., and Picard, M. (2019) Acute psychological stress increases serum circulating cell-free mitochondrial DNA. Psychoneuroendocrinology 106, 268–276

10. Yuzefovych, L. V., Pastukh, V. M., Ruchko, M. V., Simmons, J. D., Richards, W. O., and Rachek, L. I. (2019) Plasma mitochondrial DNA is elevated in obese type 2 diabetes mellitus patients and correlates positively with insulin resistance. PLoS One 14, e0222278

11. Kitazume-Taneike, R., Taneike, M., Omiya, S., Misaka, T., Nishida, K., Yamaguchi, O., Akira, S., Shattock, M. J., Sakata, Y., and Otsu, K. (2019) Ablation of Toll-like receptor 9 attenuates myocardial ischemia/reperfusion injury in mice. Biochem. Biophys. Res. Commun. 515, 442–447

12. Yang, X.-M., Cui, L., White, J., Kuck, J., Ruchko, M. V., Wilson, G. L., Alexeyev, M., Gillespie, M. N., Downey, J. M., and Cohen, M. V. (2015) Mitochondrially targeted Endonuclease III has a powerful anti-infarct effect in an in vivo rat model of myocardial ischemia/reperfusion. Basic Res. Cardiol. 110, 3

13. Mandel, P., and Metais, P. (1948) Les acides nucléiques du plasma sanguin chez l’Homme. C. R. Seances Soc. Biol. Fil. 142, 241–243

14. Bronkhorst, A. J., Ungerer, V., and Holdenrieder, S. (2019) The emerging role of cell-free DNA as a molecular marker for cancer management. Biomolecular Detection and Quantification 17, 100087

15. Tan, E. M., Schur, P. H., Carr, R. I., and Kunkel, H. G. (1966) Deoxybonucleic acid (DNA) and antibodies to DNA in the serum of patients with systemic lupus erythematosus. J. Clin. Invest. 45, 1732–1740

16. Koffler, D., Agnello, V., Winchester, R., and Kunkel, H. G. (1973) The occurrence of single-stranded DNA in the serum of patients with systemic lupus erythematosus and other diseases. J. Clin. Invest. 52, 198–204

17. Stroun, M., Anker, P., Maurice, P., Lyautey, J., Lederrey, C., and Beljanski, M. (1989) Neoplastic characteristics of the DNA found in the plasma of cancer patients. Oncology 46, 318–322

18. Fleischhacker, M., and Schmidt, B. (2007) Circulating nucleic acids (CNAs) and cancer--a survey. Biochim. Biophys. Acta 1775, 181–232

19. Sidransky, D., Von Eschenbach, A., Tsai, Y. C., Jones, P., Summerhayes, I., Marshall, F., Paul, M., Green, P., Hamilton, S. R., and Frost, P. (1991) Identification of p53 gene mutations in bladder cancers and urine samples. Science 252, 706–709

20. Wan, J. C. M., Massie, C., Garcia-Corbacho, J., Mouliere, F., Brenton, J. D., Caldas, C., Pacey, S., Baird, R., and Rosenfeld, N. (2017) Liquid biopsies come of age: towards implementation of circulating tumour DNA. Nat. Rev. Cancer 17, 223–238

21. Avriel, A., Rozenberg, D., Raviv, Y., Heimer, D., Bar-Shai, A., Gavish, R., Sheynin, J., and Douvdevani, A. (2016) Prognostic utility of admission cell-free DNA levels in patients with chronic obstructive pulmonary disease exacerbations. Int J Chron Obstruct Pulmon Dis 11, 3153–3161

22. Hummel, E. M., Hessas, E., Müller, S., Beiter, T., Fisch, M., Eibl, A., Wolf, O. T., Giebel, B., Platen, P., Kumsta, R., and Moser, D. A. (2018) Cell-free DNA release under psychosocial and physical stress conditions. Transl. Psychiatry 8, 236

23. Alunni-Fabbroni, M., Rönsch, K., Huber, T., Cyran, C. C., Seidensticker, M., Mayerle, J., Pech, M., Basu, B., Verslype, C., Benckert, J., Malfertheiner, P., and Ricke, J. (2019) Circulating DNA as prognostic biomarker in patients with advanced hepatocellular carcinoma: a translational exploratory study from the SORAMIC trial. J. Transl. Med. 17, 328

24. Perdas, E., Stawski, R., Kaczka, K., Nowak, D., and Zubrzycka, M. (2019) Altered levels of circulating nuclear and mitochondrial DNA in patients with Papillary Thyroid Cancer. Sci. Rep. 9, 14438

25. Sorber, L., Zwaenepoel, K., Deschoolmeester, V., Roeyen, G., Lardon, F., Rolfo, C., and Pauwels, P. (2017) A Comparison of Cell-Free DNA Isolation Kits: Isolation and Quantification of Cell-Free DNA in Plasma. J Mol Diagn 19, 162–168

26. Xie, J., Yang, J., and Hu, P. (2018) Correlations of Circulating Cell-Free DNA With Clinical Manifestations in Acute Myocardial Infarction. Am. J. Med. Sci. 356, 121–129

27. El Messaoudi, S., Rolet, F., Mouliere, F., and Thierry, A. R. (2013) Circulating cell free DNA: Preanalytical considerations. Clin. Chim. Acta 424, 222–230

28. Chial, H., and Craig, J. (2008) mtDNA and Mitochondrial Diseases. Nature Education 1, 217

29. Alam, T. I., Kanki, T., Muta, T., Ukaji, K., Abe, Y., Nakayama, H., Takio, K., Hamasaki, N., and Kang, D. (2003) Human mitochondrial DNA is packaged with TFAM. Nucleic Acids Res. 31, 1640–1645

30. Farge, G., and Falkenberg, M. (2019) Organization of DNA in mammalian mitochondria. Int. J. Mol. Sci. 20

31. Jorgez, C. J., Dang, D. D., Simpson, J. L., Lewis, D. E., and Bischoff, F. Z. (2006) Quantity versus quality: optimal methods for cell-free DNA isolation from plasma of pregnant women. Genet. Med. 8, 615–619

32. Thakore, N., Garber, S., Bueno, A., Qu, P., Norville, R., Villanueva, M., Chandler, D. P., Holmberg, R., and Cooney, C. G. (2018) A bench-top automated workstation for nucleic acid isolation from clinical sample types. J. Microbiol. Methods 148, 174–180

33. Stemmer, C., Beau-Faller, M., Pencreac’h, E., Guerin, E., Schneider, A., Jaqmin, D., Quoix, E., Gaub, M.-P., and Oudet, P. (2003) Use of magnetic beads for plasma cell-free DNA extraction: toward automation of plasma DNA analysis for molecular diagnostics. Clin. Chem. 49, 1953–1955

34. Trigg, R. M., Martinson, L. J., Parpart-Li, S., and Shaw, J. A. (2018) Factors that influence quality and yield of circulating-free DNA: A systematic review of the methodology literature. Heliyon 4, e00699

35. Fong, S. L., Zhang, J. T., Lim, C. K., Eu, K. W., and Liu, Y. (2009) Comparison of 7 Methods for Extracting Cell-Free DNA from Serum Samples of Colorectal Cancer Patients. Clin. Chem. 55, 587–589

36. Newell, C., Hume, S., Greenway, S. C., Podemski, L., Shearer, J., and Khan, A. (2018) Plasma-derived cell-free mitochondrial DNA: A novel non-invasive methodology to identify mitochondrial DNA haplogroups in humans. Mol. Genet. Metab. 125, 332–337

37. Mojtabanezhad Shariatpanahi, A., Rokni, P., Shahabi, E., Varshoee Tabrizi, F., and Kerachian, M. A. (2018) Simple and cost-effective laboratory methods to evaluate and validate cell-free DNA isolation. BMC Res. Notes 11, 757

38. Xue, X., Teare, M. D., Holen, I., Zhu, Y. M., and Woll, P. J. (2009) Optimizing the yield and utility of circulating cell-free DNA from plasma and serum. Clin. Chim. Acta 404, 100–104

39. Falabella, M., Kolesar, J. E., Wallace, C., de Jesus, D., Sun, L., Taguchi, Y. V., Wang, C., Wang, T., Xiang, I. M., Alder, J. K., Maheshan, R., Horne, W., Turek-Herman, J., Pagano, P. J., St Croix, C. M., Sondheimer, N., Yatsunyk, L. A., Johnson, F. B., and Kaufman, B. A. (2019) G-quadruplex dynamics contribute to regulation of mitochondrial gene expression. Sci. Rep. 9, 5605

40. Kolesar, J. E., Wang, C. Y., Taguchi, Y. V., Chou, S.-H., and Kaufman, B. A. (2013) Two-dimensional intact mitochondrial DNA agarose electrophoresis reveals the structural complexity of the mammalian mitochondrial genome. Nucleic Acids Res. 41, e58

41. Chiu, R. W. K., Chan, L. Y. S., Lam, N. Y. L., Tsui, N. B. Y., Ng, E. K. O., Rainer, T. H., and Lo, Y. M. D. (2003) Quantitative analysis of circulating mitochondrial DNA in plasma. Clin. Chem. 49, 719–726

42. Li, J., Wang, L., Yang, G., Wang, Y., Guo, C., Liu, S., Gao, Q., and Zhang, H. (2019) Changes in circulating cell-free nuclear DNA and mitochondrial DNA of patients with adolescent idiopathic scoliosis. BMC Musculoskelet. Disord. 20, 479

43. Lindqvist, D., Fernström, J., Grudet, C., Ljunggren, L., Träskman-Bendz, L., Ohlsson, L., and Westrin, Å. (2016) Increased plasma levels of circulating cell-free mitochondrial DNA in suicide attempters: associations with HPA-axis hyperactivity. Transl. Psychiatry 6, e971

44. Pearlman, S. I., Leelawong, M., Richardson, K. A., Adams, N. M., Russ, P. K., Pask, M. E., Wolfe, A. E., Wessely, C., and Haselton, F. R. (2020) Low-Resource Nucleic Acid Extraction Method Enabled by High-Gradient Magnetic Separation. ACS Appl. Mater. Interfaces 12, 12457–12467

45. Oberacker, P., Stepper, P., Bond, D. M., Höhn, S., Focken, J., Meyer, V., Schelle, L., Sugrue, V. J., Jeunen, G.-J., Moser, T., Hore, S. R., von Meyenn, F., Hipp, K., Hore, T. A., and Jurkowski, T. P. (2019) Bio-On-Magnetic-Beads (BOMB): Open platform for high-throughput nucleic acid extraction and manipulation. PLoS Biol. 17, e3000107

46. Ellison, S. L. R., English, C. A., Burns, M. J., and Keer, J. T. (2006) Routes to improving the reliability of low level DNA analysis using real-time PCR. BMC Biotechnol. 6, 33

47. Kong, F., Yuan, L., Zheng, Y. F., and Chen, W. (2012) Automatic liquid handling for life science: a critical review of the current state of the art. J Lab Autom 17, 169–185

48. Boudreau LH, Duchez A-C, Cloutier N, et al. Platelets release mitochondria serving as substrate for bactericidal group IIA-secreted phospholipase A2 to promote inflammation. Blood. 2014;124(14):2173-2183. (2015) Blood 125, 890–890

49. Xia, P., Radpour, R., Zachariah, R., Fan, A. X. C., Kohler, C., Hahn, S., Holzgreve, W., and Zhong, X. Y. (2009) Simultaneous quantitative assessment of circulating cell-free mitochondrial and nuclear DNA by multiplex real-time PCR. Genet. Mol. Biol. 32, 20–24

50. Qin, Z., Ljubimov, V. A., Zhou, C., Tong, Y., and Liang, J. (2016) Cell-free circulating tumor DNA in cancer. Chin. J. Cancer 35, 36

51. Fernando, M. R., Jiang, C., Krzyzanowski, G. D., and Ryan, W. L. (2017) New evidence that a large proportion of human blood plasma cell-free DNA is localized in exosomes. PLoS One 12, e0183915

52. Keserű, J. S., Soltész, B., Lukács, J., Márton, É., Szilágyi-Bónizs, M., Penyige, A., Póka, R., and Nagy, B. (2019) Detection of cell-free, exosomal and whole blood mitochondrial DNA copy number in plasma or whole blood of patients with serous epithelial ovarian cancer. J. Biotechnol. 298, 76–81

53. Harrington, J. S., Huh, J.-W., Schenck, E. J., Nakahira, K., Siempos, I. I., and Choi, A. M. K. (2019) Circulating mitochondrial DNA as predictor of mortality in critically ill patients: A systematic review of clinical studies. Chest 156, 1120–1136

54. Belmonte, F. R., Martin, J. L., Frescura, K., Damas, J., Pereira, F., Tarnopolsky, M. A., and Kaufman, B. A. (2016) Digital PCR methods improve detection sensitivity and measurement precision of low abundance mtDNA deletions. Sci. Rep. 6, 25186

